# Cryo-EM structures reveal bilayer remodeling during Ca^2+^ activation of a TMEM16 scramblase

**DOI:** 10.1101/420174

**Authors:** Maria E. Falzone, Jan Rheinberger, Byoung-Cheol Lee, Thasin Peyear, Linda Sasset, Ashleigh Raczkowski, Edward Eng, Annarita Di Lorenzo, Olaf S. Andersen, Crina M. Nimigean, Alessio Accardi

## Abstract

The lipid distribution of plasma membranes of eukaryotic cells is asymmetric and phospholipid scramblases disrupt this asymmetry by mediating the rapid nonselective transport of lipids down their concentration gradients. As a result, phosphatidylserine is exposed to the outer leaflet of membrane, an important step in extracellular signaling networks controlling processes such as apoptosis, blood coagulation, membrane fusion and repair. Several members of the TMEM16 family have been identified as Ca^2+^-activated scramblases but the mechanisms underlying their Ca^2+^-dependent gating and their effects on the surrounding lipid bilayer remain poorly understood. Here we describe three high-resolution cryo-electron microscopy structures of a fungal scramblase from *Aspergillus fumigatus*, afTMEM16, reconstituted in lipid nanodiscs, revealing large Ca^2+^-dependent conformational changes of the protein as well as significant, function dependent membrane reorganization.

## Introduction

The plasma membranes of eukaryotic cells are organized in an asymmetric manner; at rest, polar and charged lipids are sequestered to the inner leaflet by the activity of ATP-driven pumps. Activation of a specialized class of membrane proteins – phospholipid scramblases – causes rapid collapse of this asymmetry and externalization of negatively charged phosphatidylserine molecules. As a result, extracellular signaling networks, controlling processes such as apoptosis, blood coagulation, membrane fusion and repair, are activated (Pomorski and Menon, 2006; Nagata et al., 2016). The TMEM16 family of membrane proteins includes phospholipid scramblases and Cl^-^ channels (Falzone et al., 2018), all of which are Ca^2+^-dependent. Notably, TMEM16 scramblases also mediate Ca^2+^-dependent ion transport (Malvezzi et al., 2013; Scudieri et al., 2015; Yu et al., 2015; Lee et al., 2016; Lee et al., 2018). Prior structural and functional analyses of the fungal nhTMEM16 scramblase from *Nectria haematococca* identified a membrane-exposed hydrophilic groove that serves as the translocation pathway for ions and lipids (Brunner et al., 2014; Yu et al., 2015; Bethel and Grabe, 2016; Jiang et al., 2017; Lee et al., 2018). In the TMEM16A channel, this pathway is sealed from the membrane, thus preventing lipid access while allowing only ion permeation (Dang et al., 2017; Paulino et al., 2017a; Paulino et al., 2017b).

Phospholipid scramblases are unusual membrane proteins in the sense that their environment serves as their substrate. Activation of scramblases results in the fast and passive transbilayer movement of lipids. Thus, scramblase should remodel the membrane to create a conduit between leaflets though which lipids can diffuse. Current models of lipid translocation, based on the nhTMEM16 scramblase structure, postulate a mechanism in which lipid headgroups move through the hydrophilic permeation pathway while the tails remain embedded in the membrane core (Pomorski and Menon, 2006; Brunner et al., 2014). However, little is known about how scramblases affect the membrane structure or how Ca^2+^ binding activates TMEM16 scramblases. Here we use cryo-electron microscopy (cryo-EM) and functional experiments to address these questions. We determine the structures of the afTMEM16 scramblase reconstituted in lipid nanodiscs in an inactive (Ca^2+^-free) and an active (Ca^2+^-bound) conformation to resolutions of 3.9 Å and 4.0 Å respectively. These structures allow us to define key conformational rearrangements that underlie Ca^2+^-dependent scramblase activation. Additionally, we show that scrambling is inhibited by the lipid ceramide 24:0 (C24:0) and determine the 3.6 Å resolution structure of the Ca^2+^-bound afTMEM16/C24:0-nanodisc complex. Together, the three structures show how the scramblase interacts with its surrounding membrane substrate in different functional states.

## Results

### Structure of the Ca^2+^-bound afTMEM16 scramblase in nanodisc

To isolate conformations of afTMEM16 relevant to the scramblase activity nanodiscs were formed from a 3:1 mixture of POPE:POPG lipids, where it mediates robust lipid scrambling while its non-selective ion transport activity is silenced (Malvezzi et al., 2013; Lee et al., 2016). In the presence and absence of Ca^2+^, nanodisc-incorporated afTMEM16 adopts the TMEM16 fold (Fig. 1A, B; Fig. 1 Supplement 1-6), where each monomer in the dimeric protein comprises a cytosolic domain organized into a 6 α-helix (α1-α6)/3 β-strand (β1-β3) ferredoxin fold and a transmembrane region encompassing 10 α-helices (TM1-TM10) (Fig. 1C) (Brunner et al., 2014; Dang et al., 2017; Paulino et al., 2017a; Bushell et al., 2018). The two monomers are related by a 2-fold axis of symmetry at the dimer interface, formed by TM10 and the cytosolic domain (Fig. 1C). Five helices (TM1, TM2 and TM10 from one monomer and TM3, TM5 from the other) delimit the large and hydrophobic dimer cavity (Fig. 1D; Fig. 1 Supplement 5A), where lipids are visible in the Ca^2+^-free structure (Fig. 1 Supplement 5). In both maps the C-terminal portion of TM6 and the linker connecting it to the short cytosolic α4 helix are not well-resolved, likely reflecting their mobility within the membrane. When processing the data without imposing symmetry between subunits, the two monomers differ in the well-resolved density of the ~22 C-terminal residues of TM6 (Fig. 1 Supplement 2), likely reflecting the asymmetric orientation of the protein within the nanodisc (Fig. 1 Supplement 2A). The maps generated by signal subtracting the nanodisc and imposing C2 symmetry are of higher resolution and nearly identical to the non-symmetrized (C1) maps, apart from the resolved portions of TM6 (Fig. 1 Supplement 2, 3), and were therefore used for model building.

**Figure 1.**
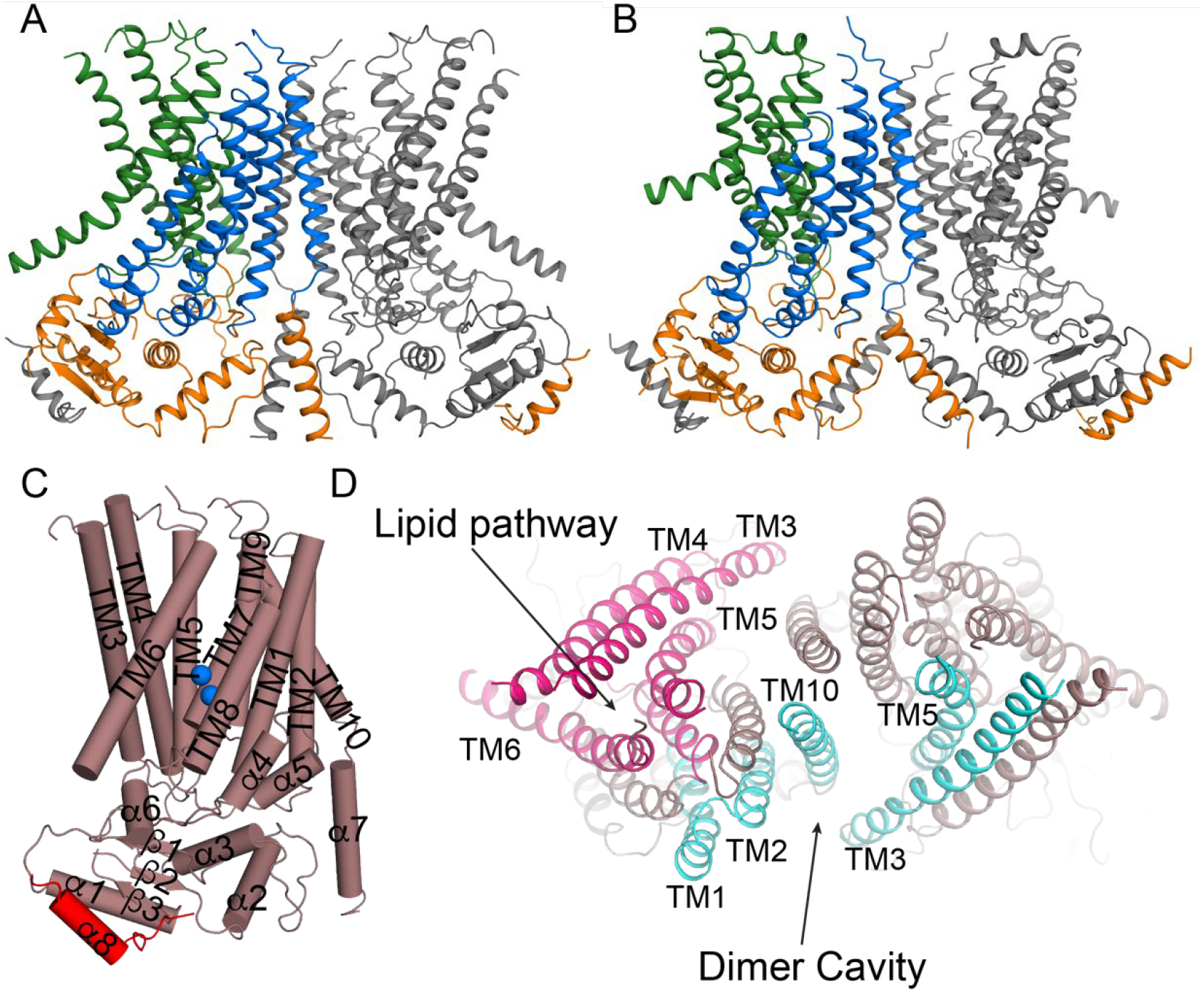
Structures of afTMEM16 in the presence and absence of Ca^2+^. (**A**) atomic models of afTMEM16 reconstituted in nanodiscs in the presence of 0.5 mM Ca^2+^ (**A**) and in the absence of Ca^2+^ (**B**). For clarity one monomer is grey, in the other the cytosolic domain is orange, the permeation pathway is green and the remainder of the protein is blue. (**C**) Helical organization of afTMEM16, the 7 domain-swapped α8 from the other monomer is shown in red and Ca2+ ions are shown as blue spheres. (**D**) Top view of Ca^2+^-bound afTMEM16 highlighting the dimer cavity (cyan) and lipid 9 translocation pathway (pink). Note that TM3 and TM5 are shared between the cavity and pathway.

In the presence of the functionally saturating concentration of 0.5 mM Ca^2+^ (Malvezzi et al., 2013), afTMEM16 adopts a conformation similar to that of Ca^2+^-bound nhTMEM16 and hTMEM16K in detergent (Brunner et al., 2014; Bushell et al., 2018), with respective r.m.s.d. ~1.8 and 2.2 Å (Fig. 1 Supplement 6). The TM3-6 from each monomer form a peripheral hydrophilic cavity that opens to the membrane with a minimum diameter of ~5 Å. This pathway allows entry and translocation of phospholipid headgroups (Fig. 1D) (Brunner et al., 2014; Yu et al., 2015; Lee et al., 2016; Jiang et al., 2017; Lee et al., 2018; Malvezzi et al., 2018) and is analogous to the ion pathway in the TMEM16A channel (Lim et al., 2016; Dang et al., 2017; Paulino et al., 2017a; Paulino et al., 2017b; Peters et al., 2018). We identified two Ca^2+^ from the density of the cryo-EM map and of an ‘omit’ difference map, calculated between experimental data and simulated maps not containing Ca^2+^ (Fig. 2A). The bound Ca^2+^ ions are coordinated by five conserved acidic residues (E445 on TM6, D511 and E514 on TM7, and E534 and D547 on TM8), three polar residues (Q438, Q518, N539), and the unpaired main chain carbonyl of G441 (Fig. 2A-C). This coordination is similar to that seen in the nhTMEM16 and hTMEM16K scramblases as well as in the TMEM16A channel, consistent with the evolutionary conservation of the Ca^2+^ binding residues (Fig. 2C) (Yu et al., 2012; Malvezzi et al., 2013; Terashima et al., 2013; Brunner et al., 2014; Tien et al., 2014; Lim et al., 2016; Dang et al., 2017; Paulino et al., 2017a; Bushell et al., 2018). In the absence of Ca^2+^ the binding site is disrupted by a movement of TM6 which displaces the three residues participating in the site (Q438, G441 and E445). Additional rearrangements of TM8, displacing N539 and E543, further disrupt the Ca^2+^ binding site (Fig. 2D-F). No Ca^2+^ density was visible in the cryo-EM map (Fig. 2E), confirming that the scramblase is in a Ca^2+^-free conformation. The movement of TM6 and TM8 in opposite directions partially relieves the electrostatic repulsion of the uncoordinated acidic side chains that form the Ca^2+^ binding site and opens a wide, water-accessible, conduit for ions to access the binding site from the intracellular solution.

**Figure 2.**
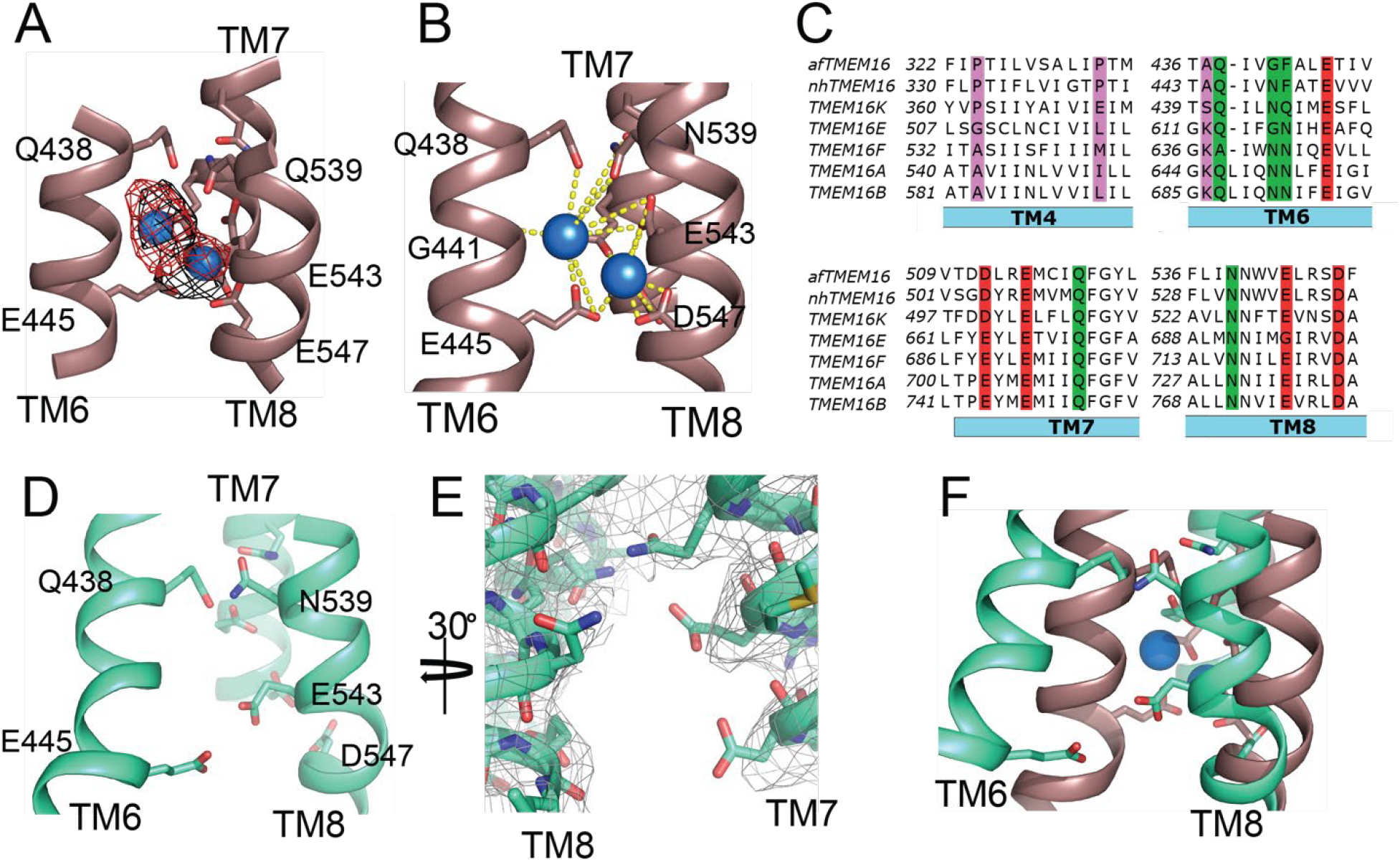
afTMEM16 Ca^2+^-binding site and conformational changes. (**A**) Close up view of the Ca^2+^-binding site, with key coordinating residues shown as sticks. The density corresponding to the Ca^2+^ ions (blue spheres) from the experimental map is shown in black and the density from the calculated omit difference map is shown in red. The peak density corresponding to the Ca^2+^ ions in the omit difference maps (red mesh) is σ=13 and 7. (**A**) Ca^2+^ coordination in afTMEM16. (**C**) Structure-based sequence alignment of the Ca^2+^-binding site and gating region of TMEM16 proteins. The alignment was generated using PROMALS3D (Pei and Grishin, 2014). Highlighted residues: conserved acidic (red) or polar (green) side chain in Ca^2+^ binding site, and the residues around which TM4 and TM6 bend (pink). (**D**) Close-up of the Ca^2+^-binding site in the absence of ligand. Coordinating residues on TM6 and TM8 are labeled. (**E**) EM density of residues lining the Ca^2+^ binding site in the absence of Ca^2+^ highlighting the lack of density for Ca^2+^ ions. (**F**) Structural alignment of the binding site with and without Ca^2+^.

### Ca^2+^ binding induces global rearrangements in afTMEM16

To understand the changes that occur upon Ca^2+^ activation, we compared our Ca^2+^-free and Ca^2+^-bound structures of afTMEM16 and identified additional conformational rearrangements of the lipid pathway and of the cytosolic domains (Fig. 3A, Fig. 3 Supplement 1). This contrasts with the TMEM16A channel, in which only the pathway-lining TM6 moves (Fig. 3B) (Paulino et al., 2017a). The cytosolic domains of afTMEM16 are translated ~3 Å parallel to the plane of the membrane, toward the axis of symmetry, such that the overall cytosolic domain becomes more compact and the cytosolic α7 helices tilt away from the axis of symmetry (Fig. 3A, right panel). In the absence of Ca^2+^, the afTMEM16 lipid pathway is closed to the membrane by a pinching motion of the extracellular portions of TM4 and TM6, which move toward each other by ~7 and ~3 Å, respectively (Fig. 3A, left panel). From the Ca^2+^-bound conformation, TM4 bends around two prolines (P324 and P333) and TM3 slides by ~6 Å (Fig. 4A) to reach the Ca^2+^-free conformation. Additionally, the intracellular portion of TM6 kinks around A437 by ~20°, inducing a ~20 Å vertical displacement of its terminal end (Fig. 4A). The displacement of TM6 is similar to that seen in the TMEM16A channel (Fig. 3B), but occurs without a glycine hinge, indicating that the movement is conserved even though the sequence is not. This leads to tighter packing of side chains from TM4 and TM6 and results in exposure of a hydrophobic surface to the membrane core (Fig. 4B). The pathway is closed to ion entry by multiple stacked aromatic and hydrophobic side chains from TM3-7 (Fig. 4C). The narrowest access point of the lipid pathway constricts from ~5 to ~1 Å preventing lipid entry (Fig. 4D-F). Residues that are important for scrambling(Jiang et al., 2017; Lee et al., 2018) cluster between helices that rearrange upon Ca^2+^ binding (Fig. 4 Supplement 1), supporting the dynamic nature of these interfaces.

**Figure 3.**
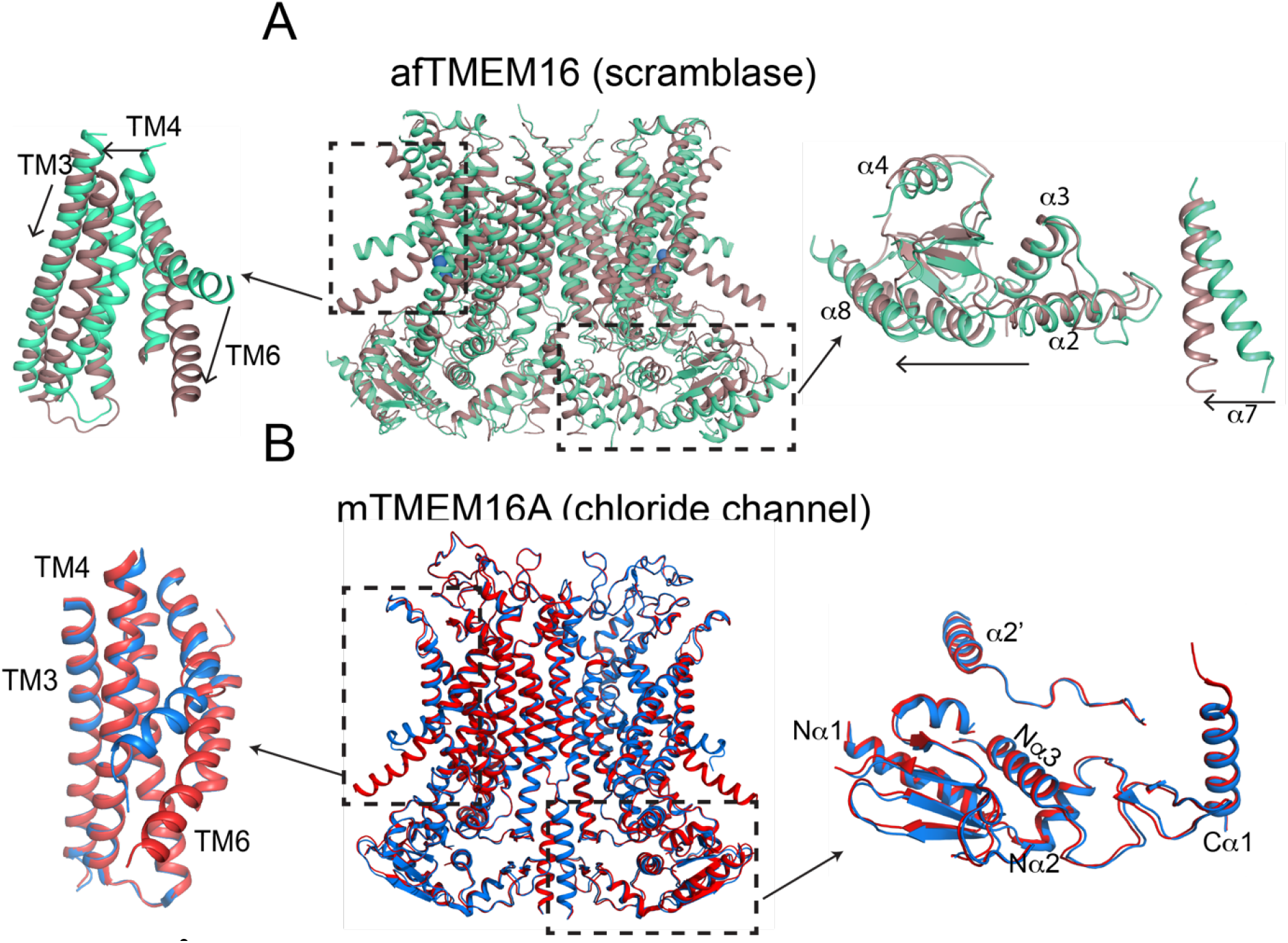
Ca^2+^-induced changes in afTMEM16 compared to TMEM16A. (**A)** Structural alignment of afTMEM16 in the presence of 0.5 mM Ca^2+^ (maroon) and absence of Ca^2+^ (cyan) (blue sphere). Left: conformational changes in lipid permeation pathway. Right: conformational changes in the cytosolic domain, Arrows indicate direction of movement from the Ca^2+^-free to the Ca^2+^-bound conformations. (**B**) Structural alignment of TMEM16A in the presence of 0.5 mM Ca^2+^ (PDBID: 5OYB, red) and absence of Ca^2+^ (PDBID: 5OYG, blue). Left: conformational changes in lipid permeation pathway. Right: view of cytosolic domains showing very a lack of movement upon Ca^2+^-binding.

**Figure 4.**
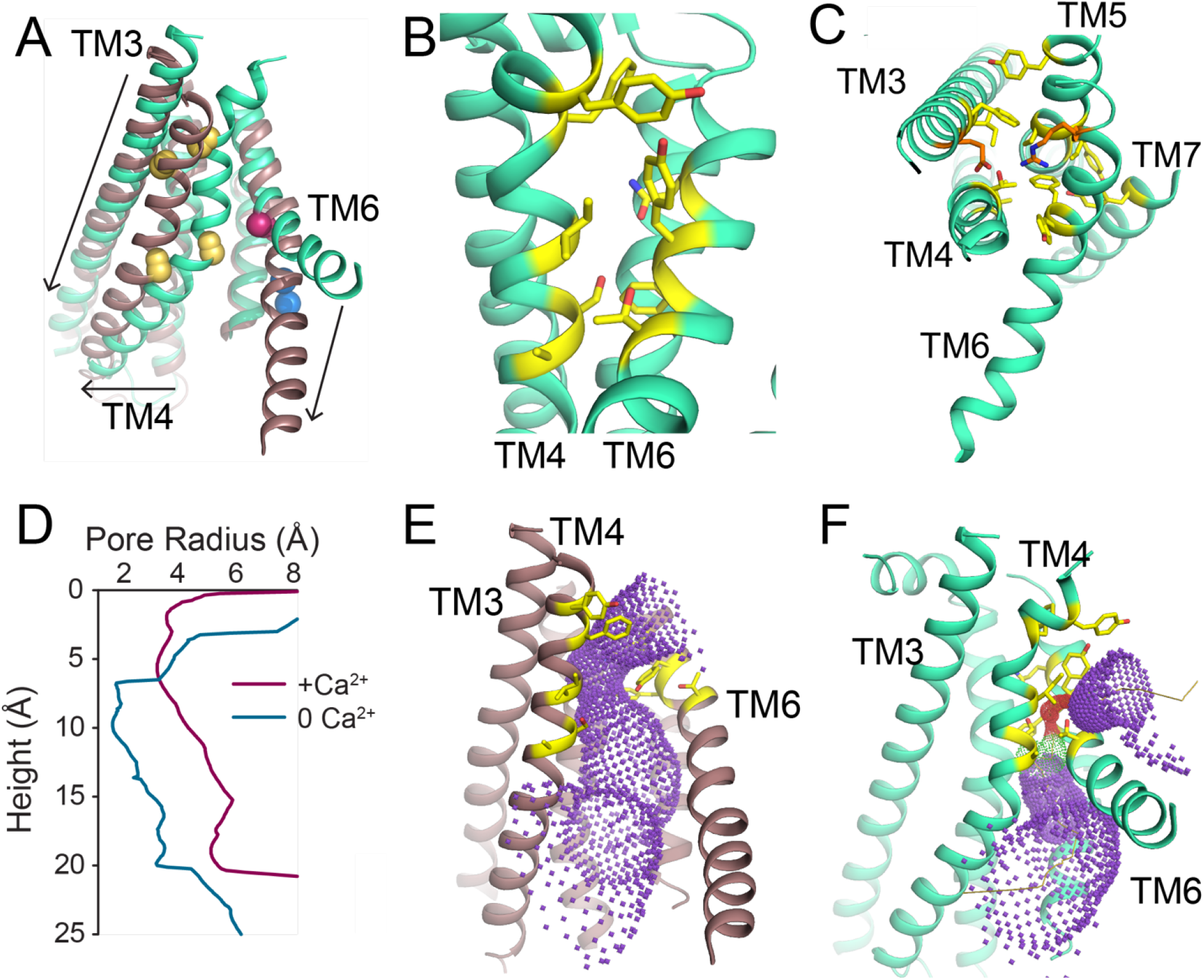
Ca^2+^-dependent rearrangements of the lipid permeation pathway. (**A**) Structural alignment of the lipid pathway with afTMEM16 in the presence (maroon) or absence (cyan) of 0.5 mM Ca^2+^. The color scheme is the same throughout the figure. Arrows indicate direction of movement from the Ca^2+^-free to the Ca^2+^-bound conformations. The lipid permeation pathway is constricted by rearrangements of TM4 around P324 and P333 (yellow spheres) and TM6 at A437 (red sphere). (**B**) Close up view of the closed permeation pathway, residues at the interface with the membrane are shown as yellow sticks. (**C**) Top view of the permeation pathway in the absence of Ca^2+^. Residues pointing inside the pathway are shown as yellow sticks. The interacting charged pair E305 and R425 are shown as orange sticks. (**D**) Diameter of the afTMEM16 lipid pathway in the presence (maroon) and absence (cyan) of Ca^2+^. The diameter was estimated using the HOLE program (Heymann, 2001; Cardone et al., 2013). (**E-F**) Accessibility of the lipid permeation pathway estimated using the program HOLE (Smart et al., 1996) in the presence (**E**) or absence (**F**) of Ca^2+^. Purple denotes areas of diameter d>5.5 Å, yellow areas where 5.5 < d < 2.75 Å and red areas with d<2.75 Å.

### Ca^2+^-dependent remodeling of the nanodisc upon scramblase activation

Because these structures are of protein/nanodisc complexes, they also provide insights into the ligand-dependent interactions between the scramblase and its substrate, the surrounding membrane. The nanodisc portion of the complex is lower in resolution than the protein, preventing us from extracting information about the detailed molecular interactions between scramblase and lipids. We nevertheless observed clear, Ca^2+^-dependent remodeling of the membrane around the protein in both the 2D class averages and 3D reconstructions (Fig. 5A-D; Fig. 5 Supplement 1). The 2D class averages are the least biased portion of our analysis to averaging or alignment artifacts, and reveal that nanodiscs containing afTMEM16 are more distorted in the presence than in the absence of Ca^2+^ (Fig. 5 Supplement 1A). This distortion is not present in 2D classes of TMEM16A-containing nanodiscs (Dang et al., 2017) or in the protein-free nanodiscs from the same sample grids (Fig. 5 Supplement 1B), suggesting that the Ca^2+^-dependent rearrangements of afTMEM16 underlie the remodeling of the nanodisc. Membrane remodeling in the presence of Ca^2+^ was observed in 3 independent datasets, whose resolutions varied between ~4 and ~9 Å for the proteinaceous regions (Fig. 5 Supplement 1C, D) and was observed independently of the processing program used for the 3D reconstructions (Fig. 5 Supplement 1E-G). Taken together these observations support the robust and reproducible nature of the observation of membrane remodeling, as well as its independence from data processing algorithms.

**Figure 5.**
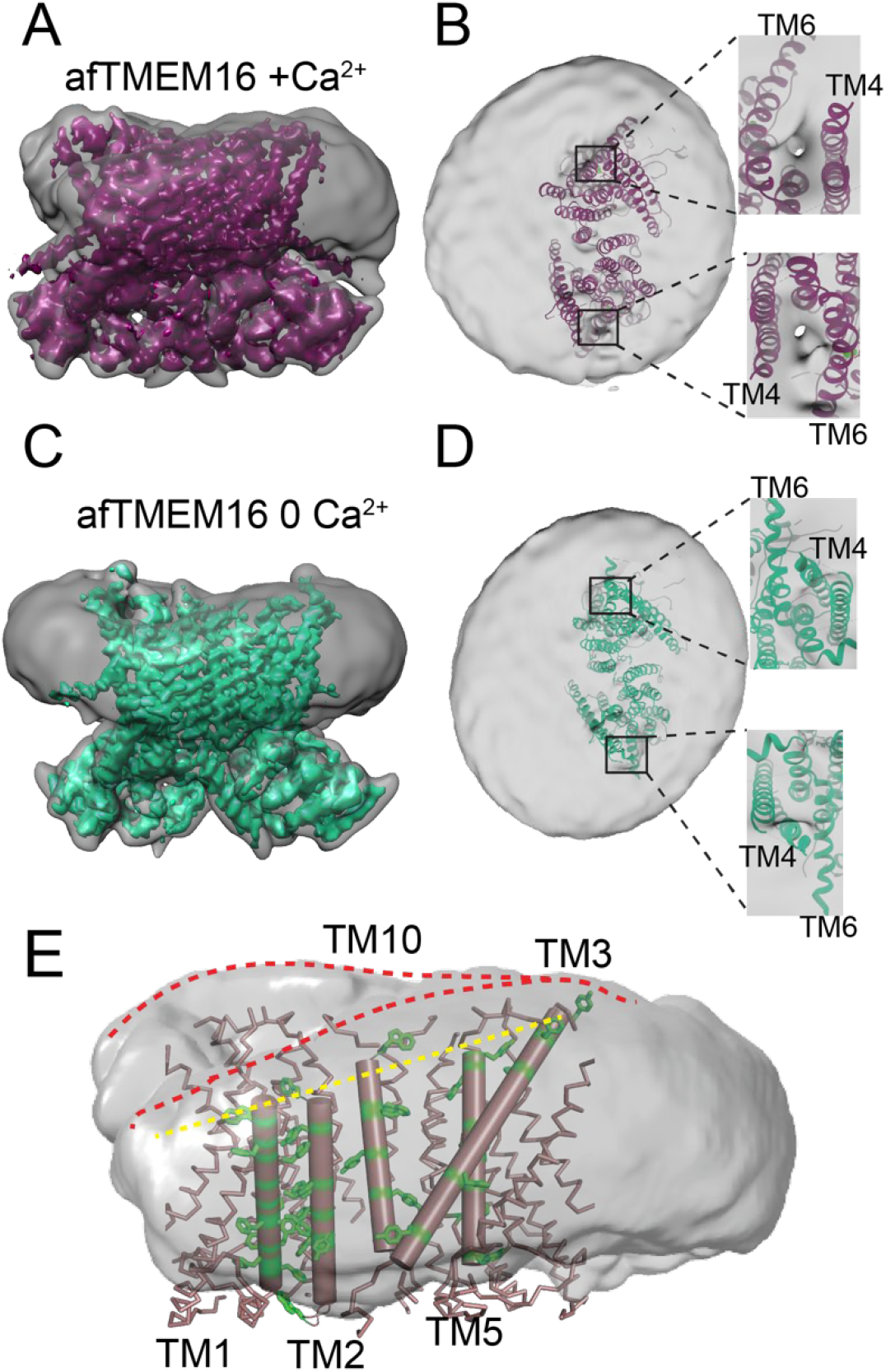
Ca^2+^-dependent membrane remodeling by afTMEM16. (**A** and **C**) masked maps of the afTMEM16/nanodisc complex with 0.5 mM Ca^2+^ (purple, **A**) and without Ca^2+^ (green, **C**) inside the respective unmasked map (gray) low pass filtered to 10 Å at σ=0.4. (**B** and **D**): top view of the unmasked map (grey) with (**B**) and without (**D**) Ca^2+^ low pass filtered to 10 Å at σ=0.1 with the respective atomic models shown as ribbons inside. Insets show close-up views of the densities at the permeation pathways. (**E**) Bending of the outer leaflet at the dimer cavity in the presence of 0.5 mM Ca^2+^. The low-pass filtered map of the complex was segmented and the isolated nanodisc density is shown. The afTMEM16 transmembrane region is shown in ribbon with the helices lining the dimer cavity (TM1, 2, 3, 5, and 10) in cartoon representation. Aromatic residues are shown as green sticks. Red dashed lines trace the upper membrane leaflet at the two sides of the lipid permeation pathway highlighting the opposite slant. Yellow dashed line traces the slope of the helices lining the dimer cavity.

The afTMEM16/nanodisc complexes are asymmetric (Fig. 5 Supplement 1H-J). In the presence of Ca^2+^ the scramblase is off-center relative to both axes of the nanodisc ellipse (Fig. 5 Supplement 1H), while in the absence of Ca^2+^ it is centered along the long but not along the short axis (Fig. 5 Supplement 1I). This raises the possibility that the acentric disposition of the protein within the nanodisc membrane depends on the activity and conformation of the protein.

3D reconstructions of the afTMEM16/nanodisc complexes with and without Ca^2+^ show that the scramblase thins the lipid bilayer near the translocation pathway and bends the outer leaflet at the dimer cavity (Fig. 5A-D). In the presence of Ca^2+^, the density of the nanodisc between the TM3-TM6 helices that form the lipid transport pathway is less than in the rest of the complex (Fig. 5B, inset). In the absence of Ca^2+^, the conformational rearrangement of TM4 and TM6 closes the pathway and no region of weak density is visible (Fig. 5D, inset). This suggests that the membrane is thinner near the lipid pathway of an active scramblase. This is consistent with the idea that the hydrophilic residues lining the open pathway provide an energetically unfavorable environment for the lipid tails, thus promoting a remodeling of the membrane (Bethel and Grabe, 2016; Jiang et al., 2017; Lee et al., 2018; Malvezzi et al., 2018). The 3D reconstructions additionally reveal that the membrane bends along the dimer cavity of afTMEM16: it is thicker around the long TM3 helix in one monomer and thinner at the short TM1 of the other (Fig. 5E). The membrane slant matches the tilted plane defined by the extracellular ends of the five cavity-lining helices (Fig. 5E). The shortest ones (TM1 and TM2) are unusually rich in aromatic side chains (Fig. 5E, Fig. 5 Supplement 2), which anchor TM segments to membrane/solution interfaces (O’Connell et al., 1990), creating a favorable environment for both phospholipid headgroups and tails. The bending of the outer leaflet along the dimer cavity is visible independent of the presence of Ca^2+^, consistent with the idea that this region does not undergo significant rearrangements upon ligand binding (Fig. 3A). This architecture is conserved in the nhTMEM16 and hTMEM16K scramblases (Fig. 5 Supplement 2A, B). However, the TMEM16A channel has a longer TM1, with fewer aromatics, and a shorter TM3 (Fig. 5 Supplement 2C, D) consistent with the lack of membrane remodeling upon its reconstitution in nanodiscs (Dang et al., 2017).

### Lipid dependence of scrambling by afTMEM16

The hypothesis that afTMEM16 remodels the membrane to scramble lipids makes two testable predictions: thicker membranes should impair scrambling and membrane remodeling should be reduced when scrambling is inhibited. Indeed, when membrane thickness was increased from ~38 to ~45 Å using lipids with longer acyl chains (Lewis and Engelman, 1983), the rate of scrambling, determined with a dithionite-based fluorescence assay(Malvezzi et al., 2013; Malvezzi et al., 2018), was reduced ~500-fold in the presence of Ca^2+^ (Fig. 6A-B, Fig. 6 Supplement 1A-D). To identify scrambling inhibitors, we considered naturally-occurring bilayer-modifying constituents of cellular membranes. In particular, we tested ceramides because they regulate cellular processes that involve activation of phospholipid scramblases, such as blood coagulation, inflammation and apoptosis (Deguchi et al., 2004; Hannun and Obeid, 2008; Borodzicz et al., 2015; Cantalupo and Di Lorenzo, 2016; Hiroshi et al., 2017). We found that addition of physiological levels of long chain ceramides potently inhibits scrambling by reconstituted afTMEM16 (Fig. 6C, Fig. 6 Supplement 1E-K). Among the tested ceramides, Ceramide 24:0 (C24:0) inhibits scrambling ~250-fold when added at 5 mole% (Fig. 6C). The inhibitory effect depends on ceramide concentration, with minimal effects at 1 mole%, and saturation, as C24:1 is nearly inert (Fig. 6C, Fig. 6 Supplement 1H). The length of the ceramide acyl chain is also important as C22:0 is as potent as C24:0 while C18:0 has minimal effect at 5 mole% (Fig. 6C, Fig. 6 Supplement 1F). The inhibitory effects of ceramides or thicker bilayers do not reflect impaired reconstitution of the protein (Fig. 6 Supplement 1L-M). We used a gramicidin-based fluorescence quench assay (Ingólfsson and Andersen, 2010) to investigate whether C24:0 and C24:1 differentially affect bulk membrane properties, such as thickness and fluidity. The assay monitors alterations in the gramicidin monomer↔dimer equilibrium, which varies with changes in membrane thickness and elasticity (Andersen et al., 2007; Lundbæk et al., 2010). Addition of C24:0 or C24:1 comparably reduces gramicidin activity (Fig. 6 Supplement 1N-P), indicating that both ceramides stiffen and/or thicken the membrane to a similar extent. The comparable effects of C24:0 and C24:1 on membrane properties suggest that their distinct effects on afTMEM16 scrambling might reflect specific interactions with the scramblase and/or their differential ability to form gel-like microdomains in membranes (Pinto et al., 2008; García-Arribas et al., 2017; Alonso and Goñi, 2018).

**Figure 6.**
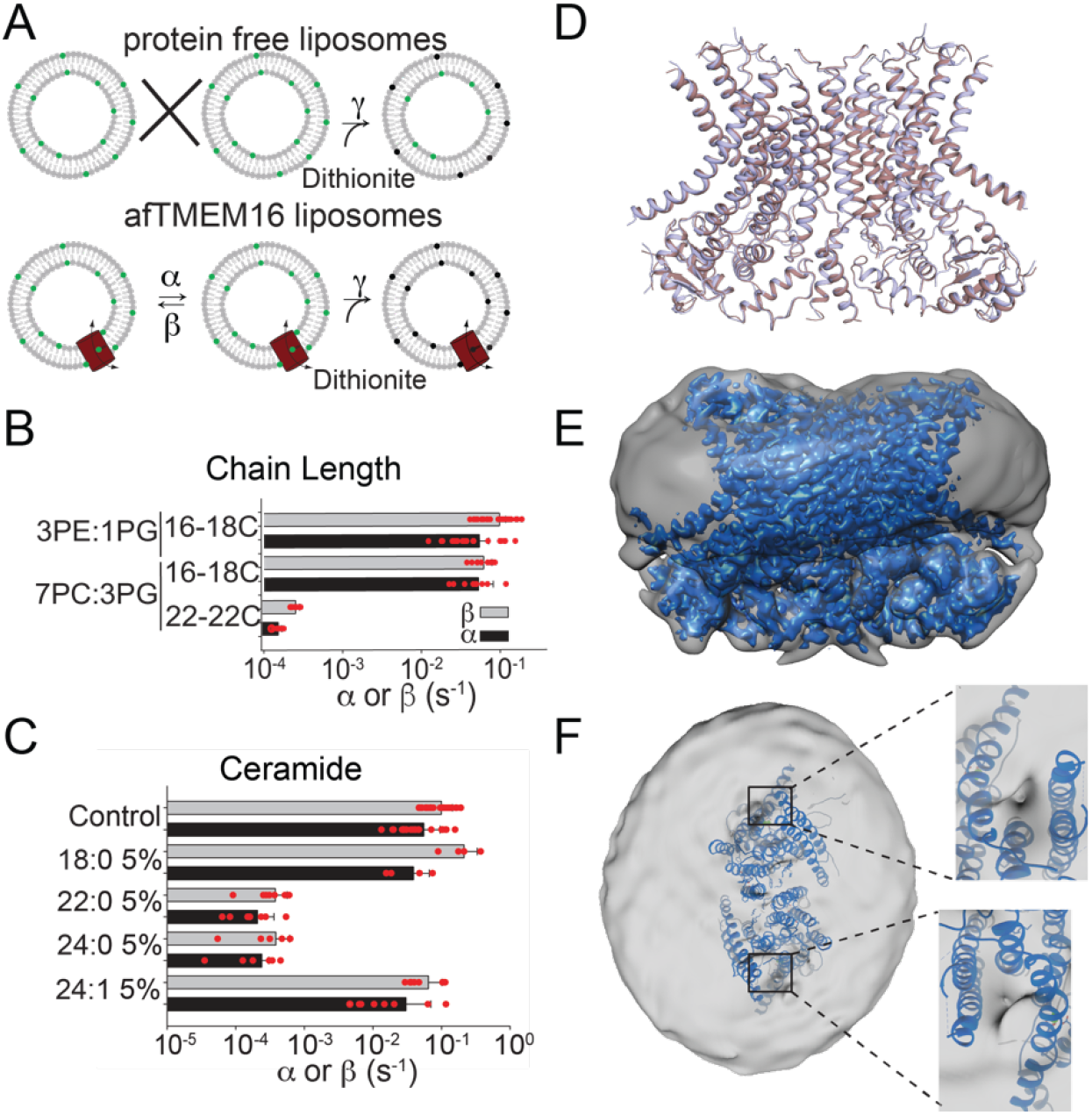
Inhibition of afTMEM16 with ceramide 24:0. (**A**) Schematic of the *in vitro* scramblase assay. Liposomes are reconstituted with NBD-labeled phospholipids (green) that distribute equally in the two leaflets. Addition of extraliposomal sodium dithionite reduces the NBD fluorophore (black), causing 50% fluorescence loss in protein free vesicles (top panel). When a scramblase is present (bottom panel), all NBD-phospholipids become exposed to dithionite, resulting in complete loss of fluorescence (Malvezzi et al., 2013). (**B-C**), Forward (α, black) and reverse (β, grey) scrambling rate constants of afTMEM16 in 0.5 mM Ca^2+^ as a function of lipid chain length (**B**) or addition of 5 mole% of different ceramides (**C**). Rate constants were determined by fitting the fluorescence decay time course to Eq. 1. A 3 POPE: 1 POPG mixture was used for the reconstitutions containing ceramide lipids (**C**). For the chain length experiments liposomes were formed from a 7 POPC: 3 POPG mixture, which does not affect the scrambling rate constants (**B**). Data is reported as mean ± St.Dev. Red circles denote individual experiments. Values and exact repeat numbers are reported in Supplementary Tables 2 and 3. (**D**) Structural alignment of Ca^2+^-bound afTMEM16 with (light blue) and without ceramide C24:0 (maroon). (**E**) Masked map of the afTMEM16/nanodisc complex in the presence of ceramide (blue) inside the unmasked map (grey) low pass filtered to 10 Å at σ=0.4. (**F**) Top view of unmasked map (grey) low pass filtered to 10 Å at σ=0.1 with the atomic models inside. Insets show the density at the permeation pathway.

### Structure of the Ca^2+^-bound and ceramide inhibited afTMEM16/nanodisc complex

To understand how ceramides affect scrambling, we determined the 3.6 Å resolution structure of Ca^2+^- bound afTMEM16 in nanodiscs containing 5 mole% C24:0 (Fig. 6D-F; Fig. 1 Supplement 1-5), a concentration that fully inhibits activity (Fig. 6C). The protein adopts a conformation where the lipid pathway is open to the membrane and both Ca^2+^-binding sites are occupied, and is nearly identical to the Ca^2+^-bound active state, with an overall r.m.s.d.<1 Å (Fig. 6D). No individual lipids are resolved within the pathway, while several partial lipid chains are visible in the dimer cavity (Fig. 1 Supplement 5C). Interestingly, although the protein adopts an active conformation, the afTMEM16/nanodisc complex is less remodeled than in the Ca^2+^-bound complex: the scramblase is centered within the nanodisc, the upper leaflet is bent at the dimer cavity (Fig. 6E) and membrane thinning at the permeation pathway is reduced (Fig. 6F). Thus, the ceramide-inhibited complex displays reduced nanodisc remodeling, even though afTMEM16 is in an open conformation. These results are consistent with the idea that phospholipid scrambling entails a thinning of the membrane near the lipid transport pathway. Finally, our findings show that while the conformation of the lipid translocation pathway is controlled by Ca^2+^ binding, the physico-chemical properties of the membrane determine whether lipid scrambling actually occurs.

## Discussion

Despite recent advances (Falzone et al., 2018), the molecular mechanisms underlying the Ca^2+^-dependent activation of the TMEM16 scramblases and their interactions with the surrounding membrane lipids remain poorly understood. Here we use cryo electron microscopy to show that the afTMEM16 scramblase reconstituted in lipid nanodiscs undergoes global conformational rearrangements upon Ca^2+^ binding (Fig. 1-3). These rearrangements involve the closure of the pathway via a pinching motion of TM4 and TM6, the upward motion of TM3 and the dilation of the Ca^2+^ binding site. The global nature of these rearrangements differs greatly from what is seen in the TMEM16A channel, where only TM6 moves in response to Ca^2+^ binding (Dang et al., 2017; Paulino et al., 2017a). Notably, similar rearrangements are observed in the human TMEM16K scramblase (Bushell et al., 2018), supporting their evolutionary conservation.

Our structural and functional experiments provide detailed insights into the activation mechanism of TMEM16 phospholipid scramblases (Fig. 1-4) and how these proteins remodel the surrounding membrane to facilitate the transfer of lipids between leaflets (Fig. 5-6). The Ca^2+^-free and Ca^2+^-bound structures of afTMEM16 define the extremes of the Ca^2+^-dependent activation process and suggest that opening of the lipid pathway is controlled by two structural elements, TM4 and TM6 (Fig. 7). Without Ca^2+^, TM4 and TM6 are bent, sealing the pathway from the lipid membrane (Fig. 7A). The first step in activation is presumably Ca^2+^ binding, which facilitates the transition of TM6 to a straight conformation and its disengagement from TM4, allowing TM6 to move towards TM8 and complete the formation of the Ca^2+^ binding site (Fig. 7B). The resulting proposed conformation is similar to that of Ca^2+^-bound TMEM16A, where TM6 is straight but lipid access is prevented by a bent TM4 (Dang et al., 2017; Paulino et al., 2017a). Straightening of the TM4 helix opens the lipid pathway to enable lipid translocation, as seen in Ca^2+^-bound afTMEM16 and nhTMEM16 (Fig. 7C)(Brunner et al., 2014). To complete the gating scheme, we propose a state where TM4 is straight and TM6 is bent (Fig. 7D), which would give rise to a partially opened lipid pathway. This conformation, while not yet observed experimentally, would account for the low, basal activity of reconstituted scramblases in the absence of Ca^2+^ (Fig. 6 Supplement 1) (Malvezzi et al., 2013; Brunner et al., 2014; Lee et al., 2016; Lee et al., 2018; Malvezzi et al., 2018). Within this gating mechanism, Ca^2+^-activated TMEM16 channels would naturally arise from scramblases via mutations that render straightening of TM4 unfavorable, while maintaining the Ca^2+^-dependent rearrangement of TM6 (Fig. 7A, B) (Dang et al., 2017; Paulino et al., 2017a). Indeed, in the Ca^2+^-free afTMEM16 scramblase, the pathway and cytosolic domain adopt conformations similar to those seen in the TMEM16A channel (Fig. 7 Supplement 1). Mutations that convert TMEM16A into a scramblase (Yu et al., 2015; Jiang et al., 2017) might re-enable the TM4 transition.

**Figure 7.**
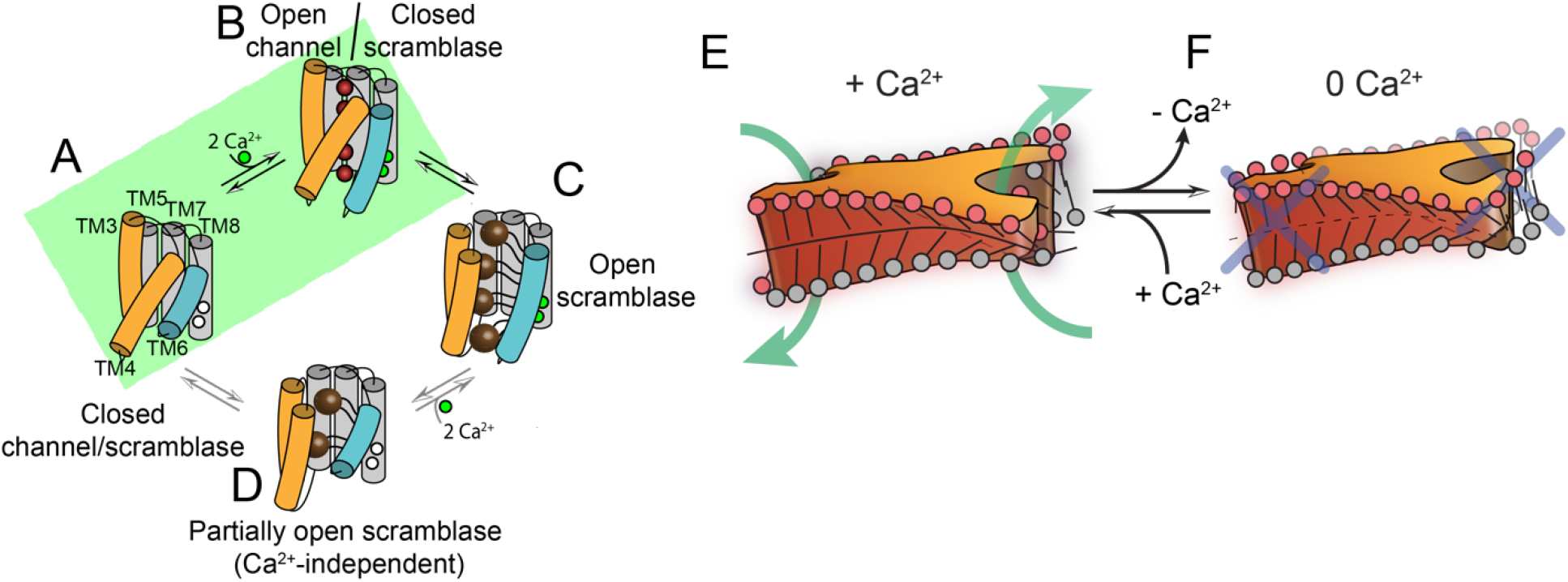
Proposed mechanisms for gating and membrane remodeling by TMEM16 scramblases. (**A-D**), Ca^2+^-dependent gating scheme for TMEM16 scramblases. The α helices lining the lipid pathway and Ca^2+^-binding site (TM3-8) are shown as cylinders. The two gating elements (TM3-4 and TM6) are respectively colored in orange and blue. When the Ca^2+^ binding sites are empty TM4 and TM6 are bent and occlude the pathway, resulting in a closed scramblase (PDBID: 6DZ7) (**A**). Upon Ca^2+^ binding, TM6 straightens and partly disengages from TM4, giving rise to a pathway that is closed to lipids but that can potentially allow ion permeation (a mTMEM16A-like state, PDBID:5OYG) (**B**). Rearrangement of TM6 promotes the straightening of TM4, resulting in an open lipid pathway (PDBID: 4WIS, 6E0H) (**C**). Straightening of TM4 in the absence of Ca^2+^, would give rise to a partially open lipid pathway that might allow the experimentally observed Ca^2+^-independent lipid scrambling (Malvezzi et al., 2013) (**D**). If the straightening of TM4 is energetically unfavorable, the scramblase is restricted to visiting only states (**A**) and (**B**), green shaded area. This would give rise to the observed Ca^2+^-dependent gating behavior of the TMEM16 channels. (**E-F**) proposed mechanism for membrane remodeling by the afTMEM16 dimer. The architecture of the dimer cavity primes the membrane by bending it in opposite directions on the two sides of the lipid translocation pathway. In the presence of Ca^2+^ the pathway is open, enabling the formation of a membrane area that is thin and forming a conduit through which lipid headgroups can translocate between the two leaflets (**E**). In the absence of Ca^2+^ the pathway is closed preventing lipid scrambling (**F**).

Our structures of afTMEM16 scramblase/nanodisc complexes suggest that lipid scrambling is the product of two features of the TMEM16 architecture: the dimer cavities and the lipid pathway (Fig. 5-6). The helices lining the dimer cavity are of different heights; TM3 and TM5 of one monomer are longer than the TM1 and TM2 from the other (Fig. 5C). The nanodisc structures suggest that the upper leaflet of the membrane bends to track the sloping cavity of the protein. A similar bending is observed in the three structures presented here (Fig. 5-6), consistent with the finding that the dimer cavity does not undergo extensive Ca^2+^-dependent conformational rearrangements. Further, while we could not identify well-defined lipids in this cavity, partial lipid densities are visible between the extracellular termini of TM10 and TM2 as well as between TM3 and TM5 in the other monomer in the Ca^2+^-free and Ca^2+^-bound/ceramide inhibited afTMEM16 structures (Fig. 1 Supplement 5). The density of the membrane near the lipid conduit is affected by the Ca^2+^-dependent conformational rearrangement of the pathway-lining helices. When the pathway is open the density is weaker, consistent with the idea that the membrane thins in this region to accommodate the remodeling of the membrane that allows the lipid headgroups to interact with the hydrophilic residues lining the lipid pathway. In the Ca^2+^-free structure, TM4 and TM6 rearrange to close the pathway preventing lipid access (Fig. 4). The membrane exposed surface of the closed pathway is hydrophobic favoring the membrane rearrangement and preventing thinning. It is important to note that these observations are robust, as a similar remodeling is seen in multiple independent datasets of varying resolutions and using different processing algorithms (Fig. 5 Supplement 1).

Based on these observations we propose a mechanism for lipid scrambling where the dimer cavity “primes” the membrane by bending the outer leaflet in opposite directions at the two sides of an open lipid pathway. This creates a membrane region that is highly curved, thin and disordered, all of which will facilitate lipid transfer between leaflets through the conduit formed by the open hydrophilic pathway (Fig. 7E) (Bennett et al., 2009; Bruckner et al., 2009; Sapay et al., 2009). In the Ca^2+^-free conformation of the scramblase, the closed pathway prevents lipid entry and membrane thinning (Fig. 7F). Similar mechanisms, where hydrophobic mismatches induce local distortion of membranes to lower the energy barrier for lipid movement through hydrophilic grooves, could be generally applicable to other scramblases. Moreover, the finding that native components of the plasma membrane, such as ceramides, specifically modulate the activity of afTMEM16 suggest that modulation of bilayer properties and composition might constitute secondary layers of regulatory control for the *in vivo* activation of scramblases.

## Methods

### Protein Expression and Purification

afTMEM16 was expressed and purified as described (Malvezzi et al., 2013). Briefly, *S. cerevisiae* carrying pDDGFP2 (Drew et al., 2008) with afTMEM16 were grown in yeast synthetic drop-out medium supplemented with Uracil (CSM-URA; MP Biomedicals) and expression was induced with 2% (w/v) galactose at 30° for 22 hours. Cells were collected, snap frozen in liquid nitrogen, lysed by cryomilling (Retsch model MM400) in liquid nitrogen (3 × 3 min, 25 Hz), and resuspended in buffer A (150 mM KCl, 10% (w/v) glycerol, 50 mM Tris-HCl, pH8) supplemented with 1 mM EDTA, 5 μg ml^-1^ leupeptin, 2 μg ml^-1^ pepstatin, 100 μM phenylmethane sulphonylfluoride and protease inhibitor cocktail tablets (Roche). Protein was extracted using 1% (w/v) digitonin (EMD biosciences) at 4°C for 2 hours and the lysate was cleared by centrifugation at 40,000 g for 45 minutes. The supernatant was supplemented with 1 mM MgCl_2_ and 10 mM Imidazole, loaded onto a column of Ni-NTA agarose resin (Qiagen), washed with buffer A + 30 mM Imidazole and 0.12% digitonin, and eluted with buffer A + 300 mM Imidazole and 0.12% digitonin. The elution was treated with Tobacco Etch Virus protease overnight to remove the His tag and then further purified on a Superdex 200 10/300 GL column equilibrated with buffer A supplemented with 0.12% digitonin (GE Lifesciences). The afTMEM16 protein peak was collected and concentrated using a 50 K_d_ molecular weight cut off concentrator (Amicon Ultra, Millipore).

### Liposome reconstitution and lipid scrambling assay

Liposomes were prepared as described (Malvezzi et al., 2013), briefly lipids in chloroform (Avanti), including 0.4% w/w tail labeled NBD-PE, were dried under N_2_, washed with pentane and resuspended at 20 mg ml^-1^ in buffer B (150 mM KCl, 50 mM HEPES pH 7.4) with 35 mM 3-[(3-cholamidopropyl)dimethylammonio]-1- propanesulfonate (CHAPS). afTMEM16 was added at 5 μg protein/mg lipids and detergent was removed using four changes of 150 mg ml^-1^ Bio-Beads SM-2 (Bio-Rad) with rotation at 4°C. Calcium or EGTA were introduced using sonicate, freeze, and thaw cycles. Liposomes were extruded through a 400 nm membrane and 20 μl were added to a final volume of 2 mL of buffer B + 0.5 mM Ca(NO_3_)_2_ or 2 mM EGTA. The fluorescence intensity of the NBD (excitation-470 nm emission-530 nm) was monitored over time with mixing in a PTI spectrophotometer and after 100 s sodium dithionite was added at a final concentration of 40 mM. Data was collected using the FelixGX 4.1.0 software at a sampling rate of 3 Hz. All experiments with added ceramide lipids were carried out in the background of 1-palmitoyl-2-oleoyl-sn-glycero-3-phosphoethanolamine (POPE), 1-palmitoyl-2-oleoyl-sn-glycero-3-phospho-(1’-rac-glycerol) (POPG) at a ratio of 3:1. Ceramides tested include: N-stearoyl-D-erythro-sphingosine (C18:0), N-behenoyl-D-erythro-sphingosine (C22:0), N-lignoceroyl-D-erythro-sphinganine (C24:0), and N-nervonoyl-D-erythro-sphingosine (C24:1) all of which were tested at 1 mole% and 5 mole%. Chain length experiments were done in the background of 7PC:3PG due to the availability of the long chain lipids. Lipids used include 1-palmitoyl-2-oleoyl-glycero-3-phosphocholine (POPC, 16-18C), POPG (16-18C), 1,2-dierucoyl-sn-glycero-3-phosphocholine (DEPC, 22-22C) and 1,2-dielaidoyl-phosphatidylglycerol (DEPG, 22-22C).

### Quantification of scrambling activity

Quantification of the scrambling rate constants by afTMEM16 was determined as recently described (Lee et al., 2018; Malvezzi et al., 2018). Briefly, the fluorescence time course was fit to the following equation

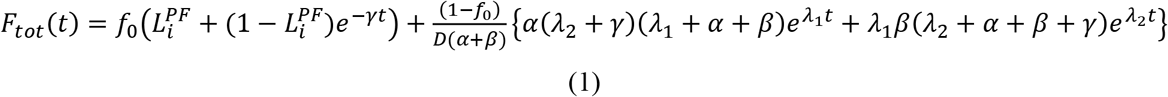

Where F_tot_(t) is the total fluorescence at time t, L_i_^PF^ is the fraction of NBD-labeled lipids in the inner leaflet of protein free liposomes, γ=γ’[D] where γ’ is the second order rate constant of dithionite reduction, [D] is the dithionite concentration, f_0_ is the fraction of protein-free liposomes in the sample, α and β are respectively the forward and backward scrambling rate constants and

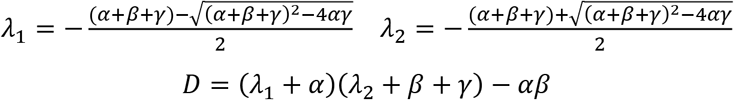

The free parameters of the fit are f_0_, α and β while L_i_^PF^ and γ are experimentally determined from experiments on protein-free liposomes. In protein-free vesicles a very slow fluorescence decay is visible likely reflecting a slow leakage of dithionite into the vesicles or the spontaneous flipping of the NBD-labeled lipids. A linear fit was used to estimate the rate of this process was estimated to be L=(5.4±1.6).10^-5^ s^-1^ (n>160). For WT afTMEM16 and most mutants the leak is >2 orders of magnitude smaller than the rate constant of protein-mediated scrambling and therefore is negligible. All conditions were tested side by side with a control preparation in standard conditions. In some rare cases this control sample behaved anomalously, judged by scrambling fit parameters outside 3 times the standard deviation of the mean for the WT. In these cases the whole batch of experiments was disregarded.

### MSP1E3 Purification and Nanodisc Reconstitution

MSP1E3 was expressed and purified as described (Ritchie et al., 2009). Briefly, MSP1E3 in a pET vector (Addgene #20064) was transformed into the BL21-Gold (DE3) strain (Stratagene). Transformed cells were grown in LB media supplemented with Kanamycin (50 mg l^-1^) to an OD_600_ of 0.8 and expression was induced with 1 mM IPTG for 3 hrs. Cells were harvested and resuspended in buffer C (40 mM Tris-HCl pH 78.0, 300 mM NaCl) supplemented with 1% Triton X-100, 5 μg ml^-1^ leupeptin, 2 μg ml^-1^ pepstatin, 100 μM phenylmethane sulphonylfluoride and protease inhibitor cocktail tablets (Roche). Cells were lysed by sonication and the lysate was cleared by centrifugation at 30,000 g for 45 min at 4° C. The lysate was incubated with Ni-NTA agarose resin for one hour at 4 °C followed by sequential washes with: buffer C + 1% triton-100, buffer C + 50 mM sodium cholate + 20 mM imidazole and buffer C + 50 mM imidazole. The protein was eluted with buffer C + 400 mM imidazole, desalted using a PD-10 desalting column (GE life science) equilibrated with buffer D (150 mM KCl, 50 mM Tris pH 8.0) supplemented with 0.5 mM EDTA. The final protein was concentrated to ~8 mg ml^-1^ (~250 μM) using a 30 kDa molecular weight cut off concentrator (Amicon Ultra, Millipore), flash frozen and stored at −80 °C.

Reconstitution of afTMEM16 in nanodiscs was carried out as follows, 3POPE:1POPG lipids in chloroform (Avanti) were dried under N_2_, washed with pentane and resuspended in buffer D and 40 mM sodium cholate (Anatrace) at a final concentration of 20 mM. Molar ratios of 1:0.8:60 MSP1E3:afTMEM16:lipids were mixed at a final lipid concentration of 7 mM and incubated at room temperature for 20 minutes. Detergent was removed via incubation with Bio-Beads SM-2 (Bio-Rad) at room temperature with agitation for two hours and then overnight with fresh Bio-Beads SM2 at a concentration of 200 mg ml^-1^. The reconstitution mixture was purified using a Superose6 Increase 10/300 GL column (GE Lifesciences) pre-equilibrated with buffer D plus 5 mM EDTA or 0.5 mM CaCl_2_ and the peak corresponding to afTMEM16-containing nanodiscs was collected for cryo electron microscopy analysis.

### Electron Microscopy Data Collection

3.5 uL of afTMEM16-containing nanodiscs (7mg mL^-1^) supplemented with 3 mM Fos-Choline-8-Fluorinated (Anatrace) was applied to a glow-discharged UltrAuFoil R1.2/1.3 300-mesh gold grid (Quantifoil) and incubated for one minute under 100% humidity at 15°C. Following incubation, grids were blotted for 2 s and plunge frozen in liquid ethane using a Vitrobot Mark IV (FEI). For the ceramide and EDTA samples, micrographs were acquired on a Titan Krios microscope (FEI) operated at 300 kV with a K2 Summit direct electron detector (Gatan), using a slid width of 20 eV on a GIFQuantum energy filter and a Cs corrector with a calibrated pixel size of 1.0961 Å /pixel. A total dose of 62.61 e^−^/Å^2^ distributed over 45 frames (1.39 e^−^/ Å^2^/frame) was used with an exposure time of 9s (200ms/frame) and defocus range of −1.5μm to −2.5μm. For the CaCl_2_ sample, micrographs were acquired on a Titan Krios microscope (FEI) operated at 300 kV with a K2 Summit direct electron detector with a calibrated pixel size of 1.07325 Å pixel. A total dose of 69.97 e^−^/Å^2^ distributed over 50 frames (1.39 e^−^/ Å^2^/frame) was used with an exposure time of 10s (200ms/frame) and a defocus range of −1.5 μm to −2.5 μm. For all samples, automated data collection was carried out using Leginon (Suloway et al., 2005).

### Image Processing

For all data sets, motion correction was performed using MotionCorr2 (Zheng et al., 2017) and contrast transfer function (CTF) estimation was performed using CTFFIND4 (Rohou and Grigorieff, 2015) both via Relion 2.0.3 (Kimanius et al., 2016). After manually picking ~2,000 particles, the resulting 2D class-averages were used as templates for automated particle picking in Relion. The particles were extracted using a box size of 275 Å with 2xbinning and subjected to 2D classification ignoring CTFs until the first peak. For the Ca^2+^-free and ceramide data sets, particles selected from 2D classification (245,835 from Ca^2+^-free, and 185,451 from ceramide) were subjected to 3D classification using the nhTMEM16 crystal structure low-pass filtered to 40 Å as an initial model. For the 0.5 mM CaCl_2_ sample two data sets were used; particles selected from the first round of 2D classification from each data set were combined and subjected to a second round of 2D classification and the resulting 302,092 particles were subjected to the same 3D classification procedure. Particles from 3D classes with defined structural features (100,268 CaCl_2_, 70,535 Ca^2+^-free, 90,709 ceramide,) were combined and re-extracted without binning and refined without enforcing symmetry using an initial model generated in CryoSPARC (Punjani et al., 2017).

Initial refinement of the afTMEM16 dataset in 0.5 mM CaCl_2_ (referred to as “+Ca^2+”^) resulted in a map with a resolution of ~7 Å. The protein was symmetric, with the exception of the resolved portion of TM6 in each monomer (Fig. 1 Supplement 2). The complex was however not two-fold symmetric due to the off-center placement of the protein within the nanodisc (Fig. 1 Supplement 2). Therefore, data processing was carried out in parallel with C2 symmetry and without enforcing symmetry (in C1 symmetry). The particles from first C1 refinement were subjected to an additional round of 3D classification, using a mask that excluded the nanodisc and maintaining the particle orientations determined by the previous refinement. The best class from 3D classification with 27,948 particles was selected for further refinement and particle polishing. Masked refinement following particle polishing resulted in a 4.36 Å final map. To refine with C2 symmetry, the particles were polished and the nanodisc was removed using signal subtraction in Relion and the subtracted particles were refined using C2 symmetry, resulting in a 4.5 Å map. Using this map, a similar procedure to the C1 processing was carried out in which the best two classes from 3D classification without alignment applying a mask including the protein (37,146 particles) were selected for further refinement. Masked refinement of these classes yielded a 4.05 Å final density map. The C1 and C2 density maps were extensively compared and determined to be nearly identical except for the resolved portion of TM6 (Fig. 1 Supplement 2). The C2 map was used for model building while the C1 map was used for analysis of the afTMEM16/nanodisc complex.

For the Ca^2+^-free data set (referred to as “0 Ca^2+^”), the first refinement resulted in a map with a resolution of ~6 Å. As with the +Ca^2+^ sample, the protein was two-fold symmetric with the exception of the resolved portion of TM6 and the overall afTMEM16/nanodisc complex was not symmetric (Fig. 1 Supplement 2). Therefore, data was processed in parallel using both C1 and C2 symmetries as described above. The C1 map was classified and the best class from 3D classification with a mask excluding the nanodisc (38,550 particles) was selected for further refinement and particle polishing. Masked refinement following particle polishing resulted in a 4.00 Å final density map. Masked, C2 refinement following polishing and signal subtraction resulted in a 3.89 Å map. The C2 map was used for model building while the C1 map was used for analysis of the afTMEM16/nanodisc complex in 0 Ca^2+^.

For the data set in the presence of 0.5 mM CaCl_2_ and 5 mole% C24:0 (referred to as “+Cera”), the selected particles were refined without symmetry, which resulted in a 4.2 Å resolution map. These particles were further classified in 3D with applied mask excluding the nanodiscs and maintaining angular information from the previous 3D refinement, from which two classes with 45,021 particles were selected for masked refinement which generated a final map of 3.74 Å. The resulting refinement showed that the nanodisc and the protein were C2 symmetric, therefore, further processing was completed with C2 symmetry enforced. Refinement resulted in a map with a resolution of 3.89 Å. An additional round of 3D classification was carried out using a mask excluding the nanodisc and maintaining the particle orientations determined by the previous refinement. The highest resolution class with 24,602 particles was selected for further refinement and particle polishing. Masked refinement following particle polishing resulted in a final map with a final average resolution of 3.59 Å.

The final resolution of all maps was determined by applying a soft mask around the protein and the gold-standard Fourier shell correlation (FSC) = 0.143 criterion using Relion Post Proccessing (Fig. 1 Supplement 1G). BlocRes from the Bsoft program was used to estimate the local resolution for all final maps (Heymann, 2001; Cardone et al., 2013)(Fig. 1 Supplement 1F). The two lower resolution +Ca^2+^ datasets mentioned were processed as described above up to the first refinement step due to the limited resolution (Fig. 5 Supplement 1C-D). For processing in cryoSPARC, extracted particles were classified in 2D and classes with structural features consistent with afTMEM16-nanodisc complexes were selected for ab initio model generation and classification. For each dataset, the best model and the associated particles were selected for homogeneous refinement followed by b-factor sharpening. Only the C24:0 dataset had a resolution better than 6 Å from cryoSPARC so these models were not used for analysis other than to confirm the observed membrane bending. For processing in cisTEM, particles from 3D classes selected in Relion were used to create an mrc stack and imported. Several rounds of automatic refinement and manual refinement with both local and global searches were carried out. Using auto-masking and manual refinement the final resolution for the +Ca^2+^ dataset was ~5 Å and was therefore not used for an analysis other than to confirm the membrane reorganization.

### Model Building, Refinement, and Validation

The maps were of high quality and we built atomic models for each structure for the transmembrane domain, most of the cytosolic region and connecting loops (Fig. 1 Supplement 4), as well as identifying seven partial lipids in the +Cera structure and two in the Ca^2+^-free structure (Fig. 1 Supplement 5). The model of afTMEM16 in the presence of ceramide was built first, due to the higher resolution of the map. The crystal structure of nhTMEM16 (PDBID 4wis) was used as a starting model and docked into the density map using chimera and the jiggle fit script in COOT (Emsley et al., 2010) (https://www2.mrc-lmb.cam.ac.uk/groups/murshudov/index.html), and mutated to match the afTMEM16 sequence. The final model contains residues 13-54, 60-119, 126-259, 270-312, 316-399, 418-461, 490-597, 602-657 and 705-724 and the following residues were truncated due to missing side chain density: L43, Q49, R53, K69, K70, E94, K100, Q102, K129, H130, D137, K257, E258, L316, E461, H555, F705, K713, E714, and E717. The model was improved iteratively by real space refinement in PHENIX imposing crystallographic symmetry and secondary structure restraints followed by manual inspection and removal of outliers. The model for the +Ca^2+^ structure was built using the refined +Cera structure (PDBID 6E1O) as an initial model, that was docked into the +Ca^2+^ density using the jigglefit script in COOT (Emsley et al., 2010) (https://www2.mrc-lmb.cam.ac.uk/groups/murshudov/index.html), manually inspected, and refined using real space refinement in PHENIX as above (Adams et al., 2010; Afonine et al., 2013). The final model contains residues 13-54, 60-119, 128-258, 270-312, 316-400, 420-460, 490-594, 602-659 and 705-724, and the following residues were truncated due to missing side chain density: D17, Q49, R53, K100, Q102, E104, K129, H130, N135, D137, E164, H247, K254, K257, H460, K550, H595, R710, K713, E714, E717, and L720. The model for the 0 Ca^2+^ structure was built using the +Cera model as a starting point and regions where the density differed were built manually (TM3, TM4 and TM6) or rigid body refined where smaller rearrangements were observed. The final model contains residues 13-54, 60-119, 128-258, 270-312, 316-400, 420-460, 490-594, 602-659 and 705-724, and the following residues were truncated due to missing side chain density: D17, F36, R53, K70, E94, K100, Q102, E132, K241, H247, R250, K257, E310, K317, K634, K642, Y654, R704, R708, E714, E717. For all three structures, the density in the regions of TM3 and TM4 and the connecting loop (residues 274-352) was less well-defined than remainder of the structure and the model may be less accurate in this area.

To validate the refinement, the FSC between the refined model and the final map was calculated (FSCsum) (Fig. 1 Supplement 1H). To evaluate for over-fitting, random shifts of up to 0.3 Å were introduced in the final model and the modified model was refined using PHENIX against one of the two unfiltered half maps.. The FSC between this modified-refined model and the half map used in refinement (FSCwork) was determined and compared to the FSC between the modified-refined model and the other half map (FSCfree) which was not used in validation or refinement. The similarity in these curves indicates that the model was not over-fit (Fig. 1 Supplement 1H). The quality of all three models was assessed using MolProbity (Chen et al., 2010) and EMRinger (Barad et al., 2015), both of which indicate that the models are of high quality (Fig. 1 Supplement 1H).

### Difference map calculation

To compare the maps resulting from C1 and C2 processing in the 0 Ca^2+^ and +Ca^2+^ structures, we calculated a difference map between the two volumes using the volume operation subtract function in chimera (Pettersen et al., 2004) (Fig. 1 Supplement 2). We used ‘omit density’ to assign the placement of several lipids in the 0 Ca^2+^ and +Cera structures and the Ca^2+^ ions in the +Ca^2+^ and +Cera structures which were calculated using the ‘phenix.real_space_diff_map’ function in PHENIX (Adams et al., 2010; Afonine et al., 2013; Dang et al., 2017). Briefly, completed models without the ligand in question were used to generate a theoretical density map which was subtracted from the experimental density map. In the subtracted map, areas of ligand density appeared as positive density. (Schneider et al., 2012)

### Gramicidin Fluorescence Assay

The gramicidin channel-based fluorescence assay for monitoring changes in lipid bilayer properties was implemented using large unilamellar vesicles (LUVs) incorporating gramicidin (gA) and loaded with 8-aminonaphthalene-1,3,6-trisulfonic acid, disodium salt (ANTS), following published protocols (Ingólfsson and Andersen, 2010; Ingólfsson et al., 2011). Vesicles composed of 1-palmitoyl-2-oleoyl-sn-glycero-3-phosphoethanolamine (POPE) and 1-palmitoyl-2-oleoyl-sn-glycero-3-phospho-1’-rac-glycerol (POPG) in 3:1 (mol/mol) proportion in chloroform were mixed with a gA to lipid molar ratio of 1:40,000. An additional 0, 1, or 5 mole-percent of C24 ceramide (d18:1/24:0) or C24:1 ceramide (d18:1/24:1) in chloroform were added to the gA-phospholipid mix. The lipid mixtures were dried under nitrogen to remove chloroform and further dried in vacuum overnight. Lipids were rehydrated with fluorophore-containing buffer solution (25 mM ANTS, 100 mM NaNO_3_, and 10 mM HEPES), vortexed for 1 min, and incubated at room temperature for at least 3 hours while shielded from light. The solutions were sonicated for 1 min at low power and subjected to 6 freeze-thaw cycles. The samples then were extruded 21 times with an Avanti Polar Lipids mini-extruder (Avanti) and 0.1 mm polycarbonate filter. The samples were extruded at 35-40°C. Extravesicular ANTS was removed with PD-10 desalting column (GE Healthcare,). The average LUV diameter was ~130 nm, with an average polydispersity index (PDI) of 10%, as determined using dynamic light scattering. LUV stock solutions were stored at room temperature with an average shelf life of 7 days.

The time course of the ANTS fluorescence was measured at 25 °C with an SX-20 stopped-flow spectrofluorometer (Applied Photophysics), with an LED light source. Excitation was at 352 nm and emission was recorded above 450 nm with a high pass filter; the deadtime of the instrument is ≈ 2 ms with a sampling rate of 5000 points/s. Samples were prepared by diluting the stock lipid concentration with buffer solution (140 mM NaOH and 10 mM HEPES) to 125 μM LUV; each sample was incubated for at least 10 min before several 1 s mixing reactions. Each sample was first mixed with the control buffer (no Tl^+^), followed by mixing with the quench solution (50 mM TINO_3_ 94 mM NaNO_3_ and 10 mM HEPES). Experiments are conducted with 2 independently prepared populations of vesicles per lipid/ceramide combination and traces are analyzed in MATLAB (MathWorks,).

The fluorescence quench rate therefore was determined as described (Ingólfsson and Andersen, 2010) by fitting the time course to a stretched exponential function (Berberan-Santos et al., 2005):

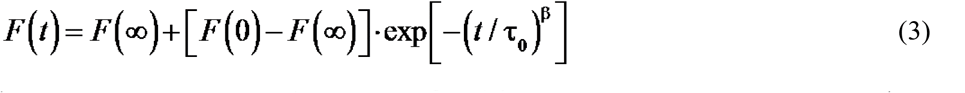

*F*(*t*) denotes the fluorescence intensities at time *t*; β (0 < β ≤ 1) is a parameter that accounts for the dispersity of the vesicle population; and τ_o_ is a parameter with dimension of time. *F*(0), *F*(∞), β and τ_o_ were determined from a nonlinear least squares fit of Eq. 1 to each individual fluorescence trace, and the quench rate was determined (Berberan-Santos et al., 2005):

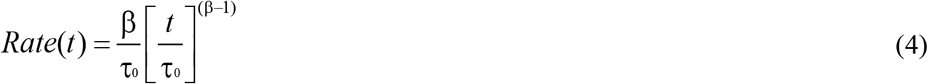

evaluated at *t* = 2 ms. The ceramide-induced changes in the quench rate then was evaluated as

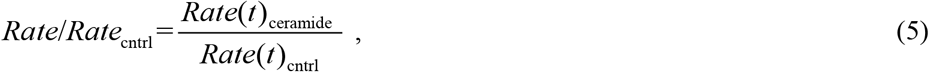

where the subscripts “ceramide” and “cntrl” denote the rates observed in the presence and absence of ceramide.

### Data Availability

The three-dimensional cryo-EM density maps of the calcium-bound, calcium-free, and Ca^2+^-bound in the presence of ceramide 24:0 afTMEM16/nanodisc complexes have been deposited in the Electron Microscopy Data Bank under accession numbers EMD-8948, EMD-8931, and EMDB-8959 respectively. The deposition includes corresponding masked and unmasked maps, the mask used for the final FSC calculation, and the FSC curves. Coordinates for the models of the calcium-bound, calcium-free, and Ca^2+^-bound ceramide inhibited states have been deposited in the Protein Data Bank under accession numbers 6E0H, 6DZ7, and 6E1O respectively. All other data are available from the corresponding author upon reasonable request.

## Author contributions

M.E.F. and A.A. designed research; M.E.F., A.R. and E.E. performed cryo-electron microscopy data collection; M.E.F. and J.R. analyzed cryo-EM data; M.E.F. and B.-C.L. performed and analyzed TMEM16 scrambling experiments; A.D.L and L.S. assisted with ceramide experiments; O.S.A. and T.P. performed and analyzed gramicidin flux assays; M.E.F. and A.A. wrote the manuscript; M.E.F., C.M.N., O.S.A., J.R. and A.A. edited the manuscript.

## Acknowledgements

The authors thank members of the Accardi lab for helpful discussions. Richard Hite for helpful suggestions on cryo-EM data processing, Christopher Miller and Simon Scheuring for insightful discussions on the work, Jeremy Dittman for insightful discussions and artistic contributions to the work, Eva Fortea Verdejo for help with sequence alignments and artistic contributions, Olga Boudker and Xiaoyu Wang for help with the nanodisc preparation. This work was supported by NIH Grant R01GM106717 (to A.A.) and 1R01GM124451-02 (to C.M.N.), an Irma T. Hirschl/Monique Weill-Caulier Scholar Award (to A.A.), by the Basic Science Research Program through the National Research Foundation of Korea (N.R.F.) funded by the Ministry of Education, Science and Technology (grant 2013R1A6A3A03064407 to B.-C. L.). M.E.F. is the recipient of a Weill Cornell Medicine Margaret & Herman Sokol Fellowship. All EM data collection and screening was performed at the Simons Electron Microscopy Center and National Resource for Automated Molecular Microscopy located at the New York Structural Biology Center, supported by grants from the Simons Foundation (349247), NYSTAR, and the NIH National Institute of General Medical Sciences (GM103310) with additional support from Agouron Institute (F00316) and NIH S10 OD019994-01. Initial negative stain screening was performed at the Weill Cornell Microscopy and Image Analysis Core Facility, with the help of L. Cohen-Gould, and at the Rockefeller University Electron Microscopy Resource Center, with the help of Kunihiro Uryu and Devrim Acehan.

---

**Figure 1-Supplement 1.**
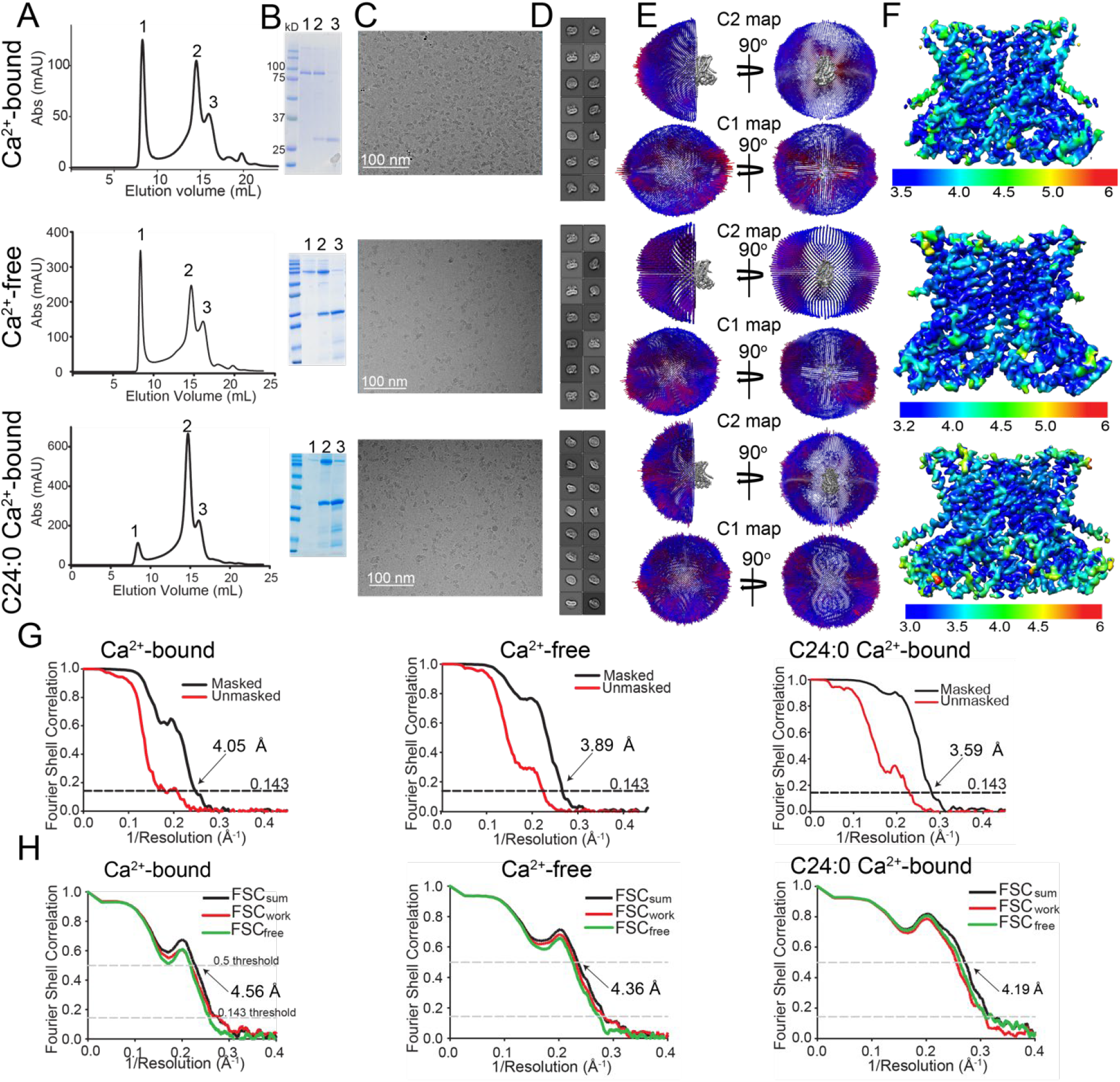
Cryo-EM characterization of afTMEM16/nanodisc. (**A-F**) *Top:* + 0.5 mM Ca^2+^. *Middle*: 0 Ca^2+^. *Bottom*: + 0.5 mM Ca^2+^ and + 5 mole% C24:0. (**A**) Size exclusion profile of afTMEM16 in MSP1E3 nanodiscs. (**B**) SDS gel of the indicated size exclusion peaks using precision plus kaleidoscope ladder. Molecular weight (MW) of afTMEM16 is 84.5 kD and MW of MSP1E3 is 32.5 kD. (**C**) Representative cryo-EM micrographs of vitrified afTMEM16/nanodisc complexes. (**D**) Representative 2D-class averages, box size 275 Å. (**E**) Angular distribution representation of final, signal-subtracted C2 or C1 maps, number of views at each angular orientation is represented by length and color of cylinders where red indicates more views. (**F**) Final masked reconstruction colored by local resolution calculated using the Bsoft program BlocRes (Heymann, 2001; Cardone et al., 2013). (**G**) FSC plot indicating the resolution at the 0.143 threshold of final masked (black) and unmasked (red) map of afTMEM16 in the presence of 0.5 mM Ca^2+^ (right) absence of Ca^2+^ (middle), and presence of 0.5 mM Ca^2+^ and 5 mole% C24:0. (**H**) FSC curves of refined models versus maps of afTMEM16 reconstituted in nanodiscs in the presence of 0.5 mM Ca^2+^ (right), in the absence of Ca^2+^ (middle) or with 0.5 mM Ca^2+^ and 5 mole% C24:0 ceramide (right). Black traces: FSC curves for the refined model compared to the final masked reconstruction (FSC_sum_). Red traces: FSC curves for the modified, refined model compared to the masked half-map 1 (FSC_work_, used during validation refinement). Green traces: FSC curves for the modified, refined model compared to the masked half-map 2 (FSC_free_, not used during validation refinement). Dashed lines show FSC threshold used for FSC_sum_ of 0.5 and for FSC_free/work_ of 0.143. Statistics for the EM analysis and model building are reported in Supplementary Table 1.

**Figure 1-Supplement 2.**
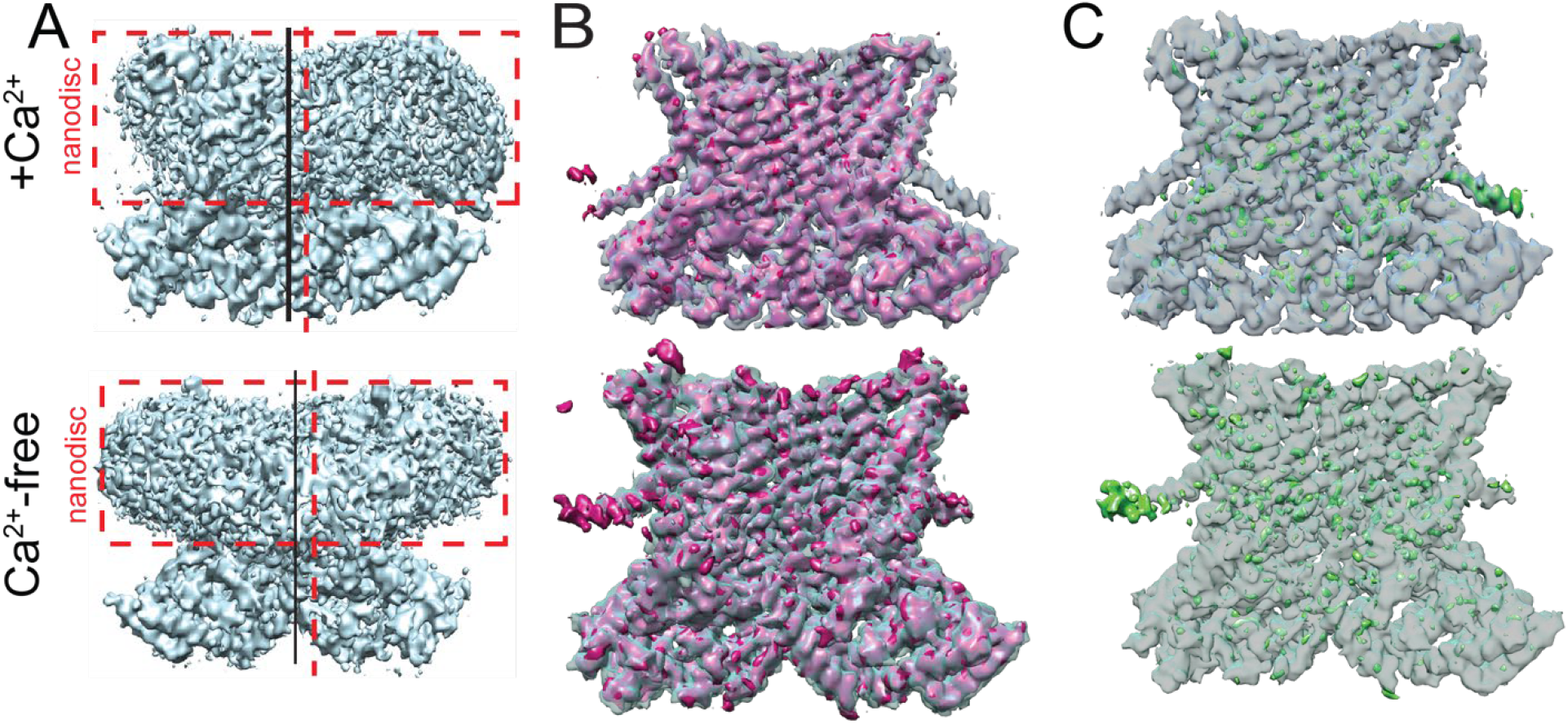
Asymmetry of afTMEM16-nanodisc complex. (**A**) Unmasked final C1 map highlighting the non-centered positioning of the protein within the nanodisc in the presence (top) and absence (bottom) of Ca^2+^. Dashed red box is drawn around the nanodisc and dashed vertical red line shows the symmetry axis of the nanodisc. Vertical black line shows the symmetry axis of the protein. (**B**) Overlay of C1 (solid pink) and C2 (transparent gray) maps for the + Ca^2+^ (top) and 0 Ca^2+^ complexes (bottom). (**C**) The difference map between the C1 and C2 maps (green) is shown inside the C2 map (gray transparent).

**Figure 1-Supplement 3.**
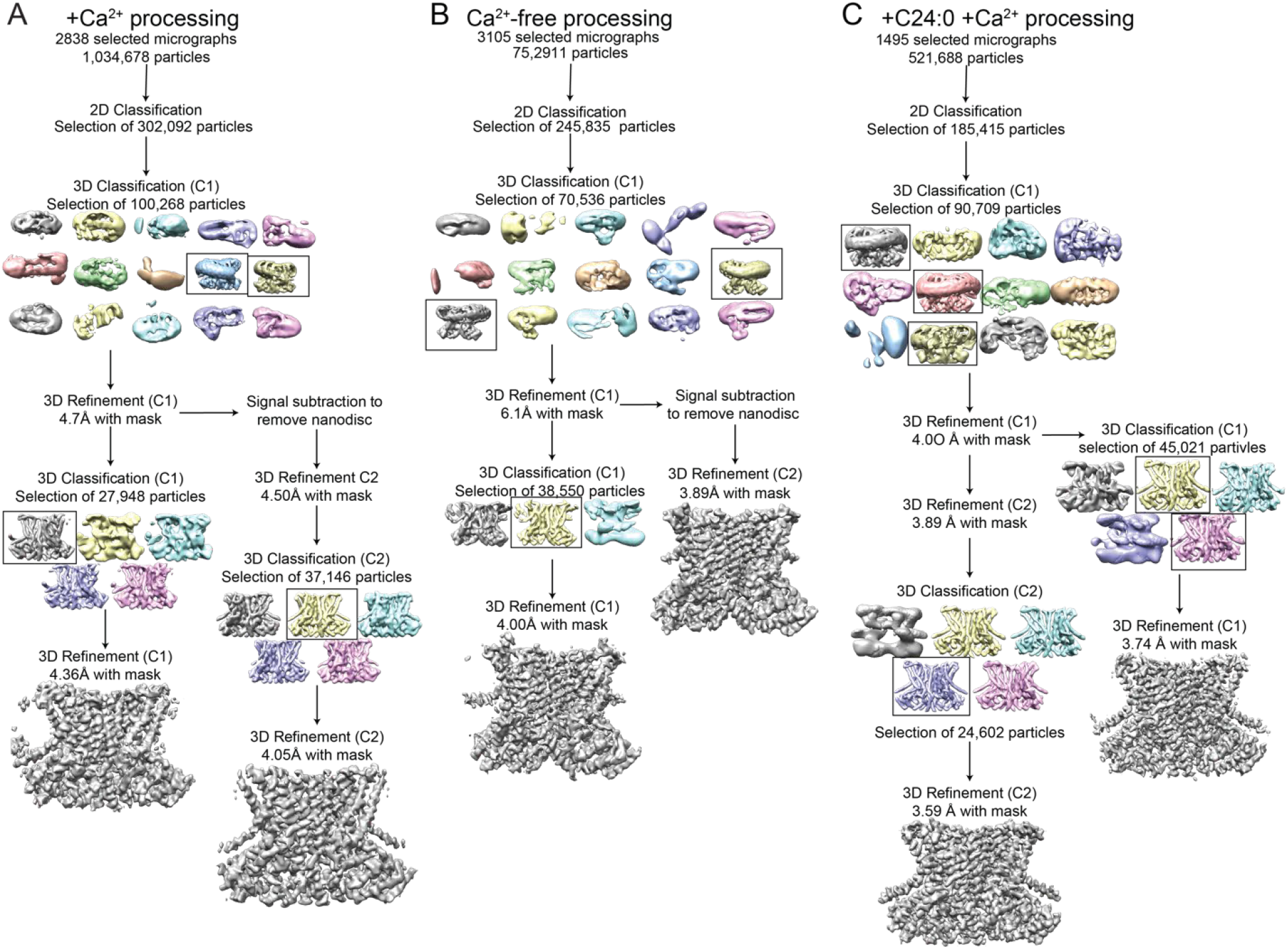
Cryo-EM data processing procedure for afTMEM16/nanodisc complexes with and without Ca^2+^. (**A-C**) Data processing flow chart for +Ca^2+^ (**A**), Ca^2+^-free (**B**), and +C24:0 +Ca^2+^ (**C**). Particles picked from manually inspected micrographs were sorted with 2D and 3D classification in C1 before particle polishing and final masked refinement. For the +Ca^2+^ and Ca^2+^-free structures, particles were further classified and refined in C2 following signal subtraction of the nanodiscs density. See methods for detailed procedure.

**Figure 1-Supplement 4.**
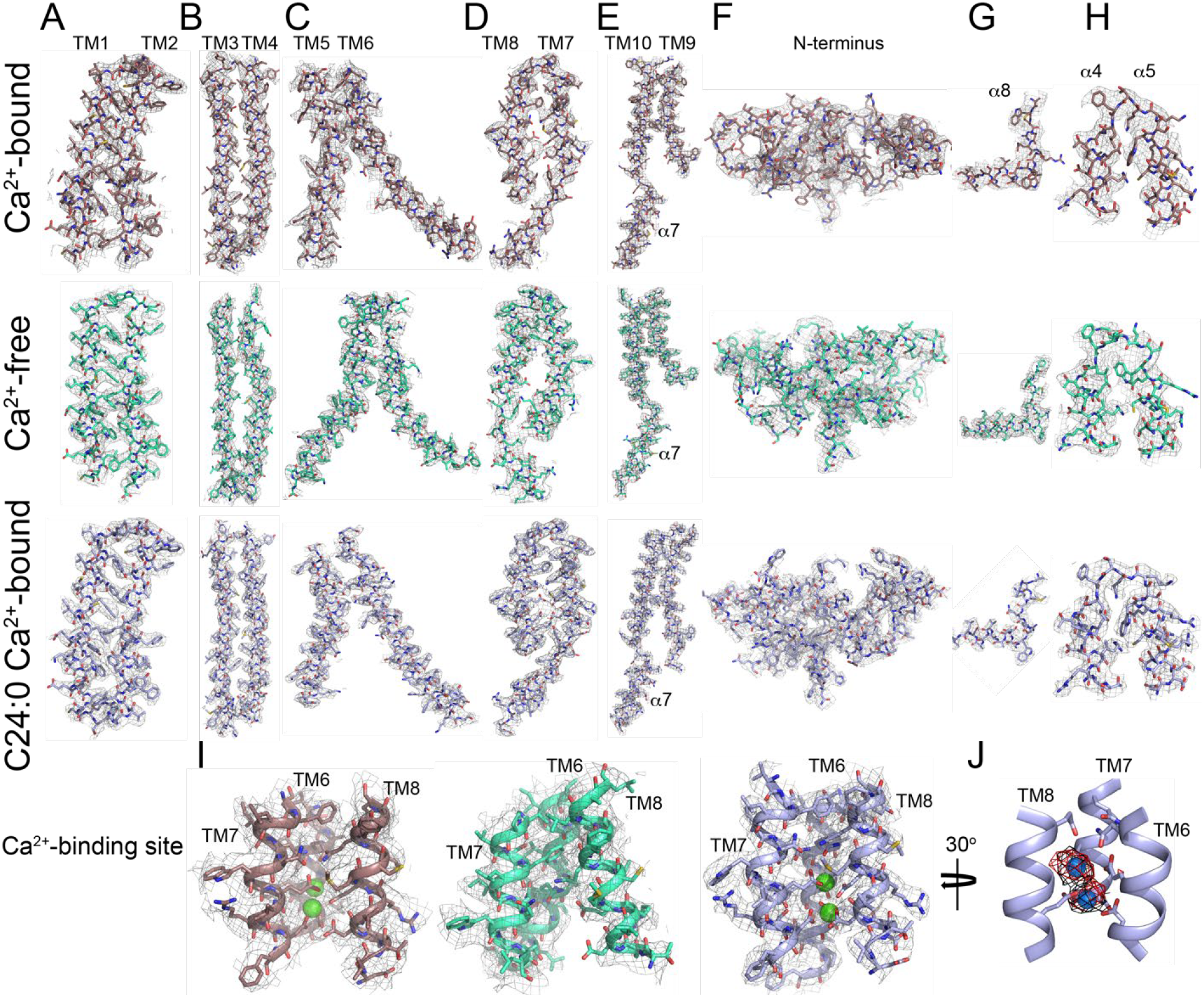
Representative cryo-EM density for afTMEM16 with and without Ca^2+^. (**A**-**I**), The cryo-EM density (gray mesh) is overlaid with the respective atomic models (sticks), Ca^2+^-bound afTMEM16 is in maroon (top panels), Ca^2+^-free is in cyan (middle panels) and Ca^2+^-bound afTMEM16 in the presence of C24:0 is in light blue (bottom panels). Heteroatoms are colored as follows: oxygen is red, nitrogen is blue, and sulfur is yellow. (**A-E**), transmembrane domains TM1-10 and cytosolic α7; (**F**) N-terminus containing *α*1-3, α6 and β1-3. (**G**) Close up of the domain-swapped region α8. (**H**) Close up of the cytosolic helices α4-5. (**I**) Ca^2+^ binding site consisting of portions of TM6-8 with Ca^2+^ ions shown in green. (**J**) Placement of Ca^2+^ ions (green) in the +Ca^2+^ + C24:0 structure inside EM density map (black) and omit difference map (red).

**Figure 1-Supplement 5.**
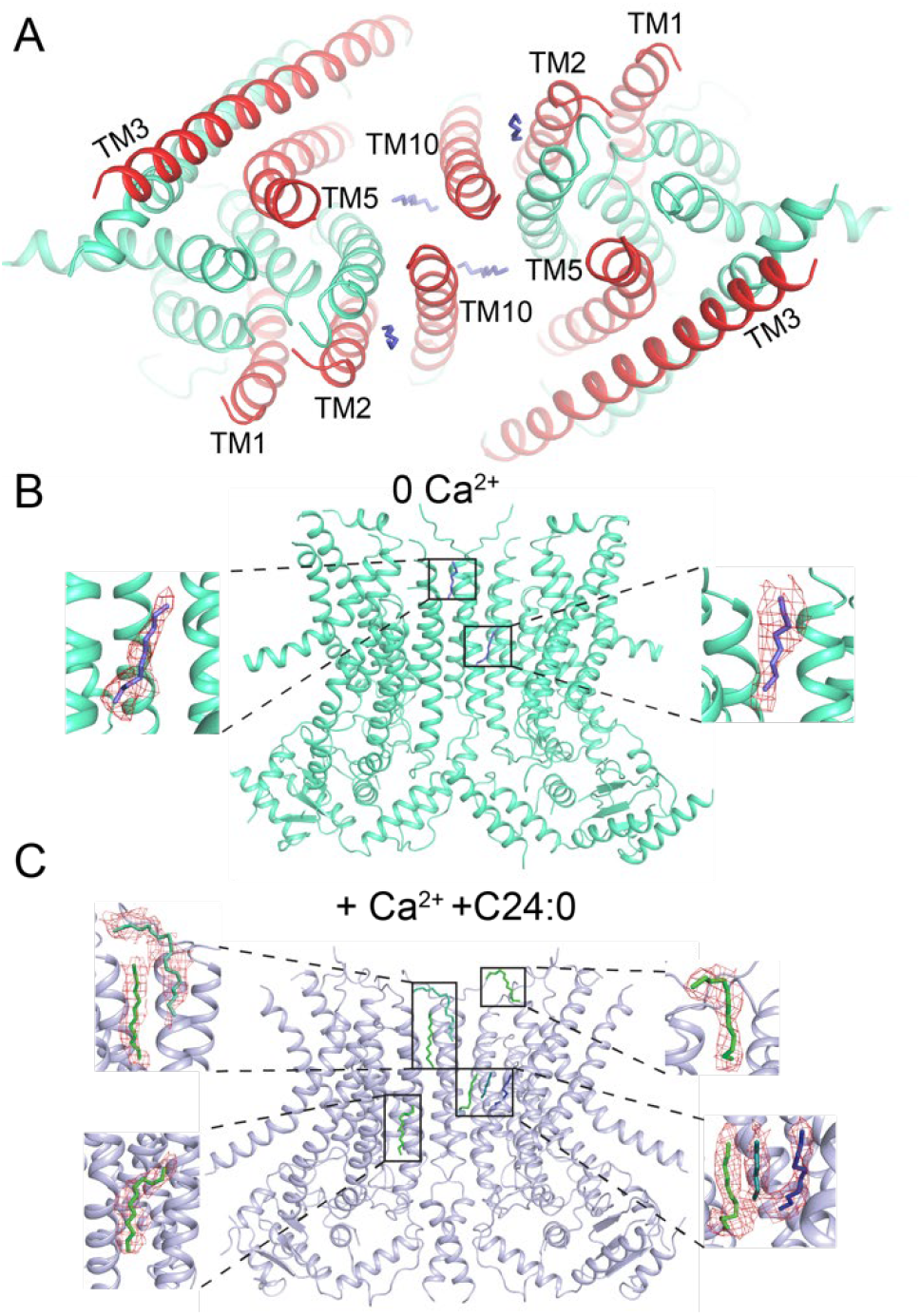
Lipids in dimer cavity in the absence of Ca^2+^. (**A-B**) Top (**A**) or side view (**B**) of the atomic model of afTMEM16 in the absence of Ca^2+^ with location of lipid acyl chains (in purple) within the dimer cavity. (**C**) Side view of the atomic model of afTMEM16 in the presence of 0.5 mM Ca^2+^ and 5 mole% C24:0 with location of lipid acyl chains within the side dimer cavity. Insets show acyl chains colored by ligand ID (D12-green, D10-blue, 8K6-cyan, OCT-teal) into the difference map density (in red) between the experimental density map and the theoretical map calculated from the model with lipids omitted. Only the lipids in one dimer cavity are shown for clarity.

**Figure 1-Supplement 6.**
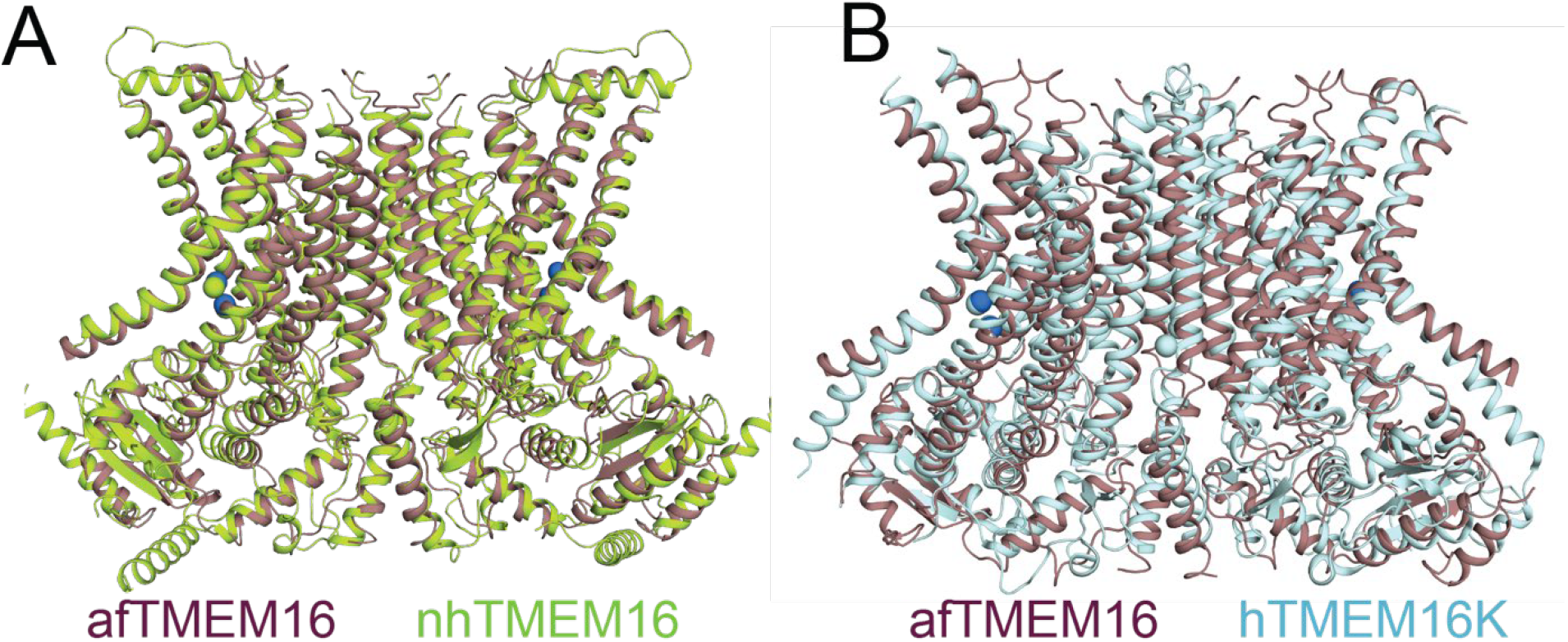
afTMEM16 in nanodiscs is very similar to nhTMEM16 and human TMEM16K in detergent. (**A-B**) Structural alignment of the atomic models of Ca^2+^-bound afTMEM16 in nanodiscs (purple) and nhTMEM16 in detergent (limon, PDBID 4WIS) of hTMEM16K in detergent (light blue, PDBID 5OC9).

**Figure 3-Supplement 1.**
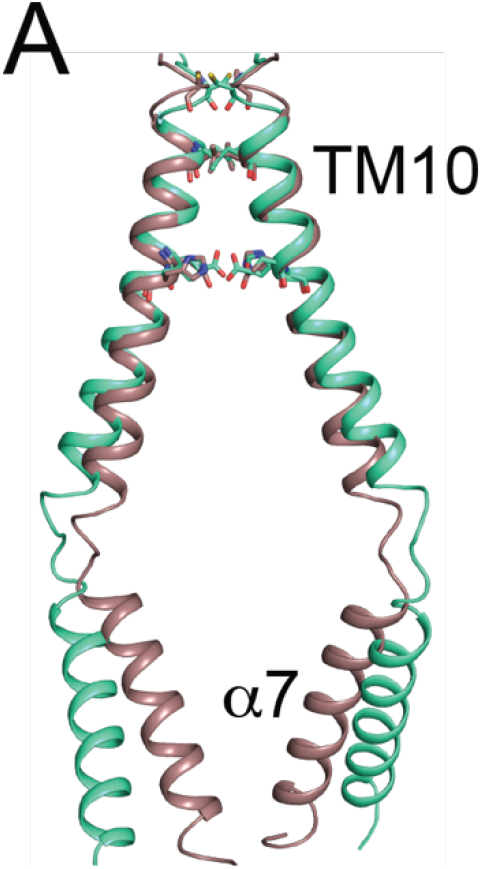
afTMEM16 dimer interface. (**A**)Structural alignment of the TM10-α7 dimer interface of afTMEM16 in Ca^2+^-bound (maroon) and Ca^2+^-free (cyan) conformations.

**Figure 4-Supplement 1.**
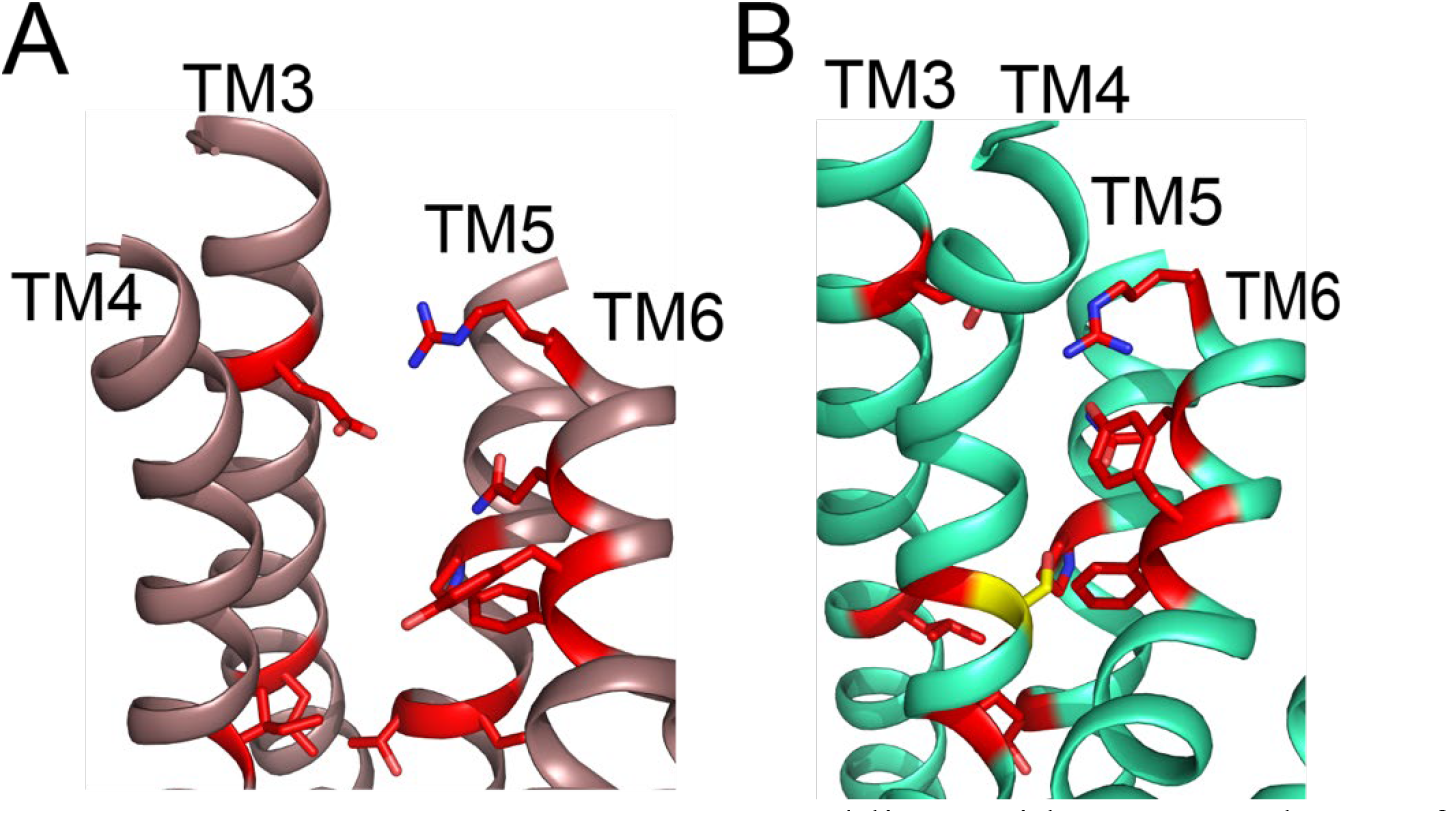
Important scrambling residues mapped onto afTMEM16 open and close permeation pathways. (**A-B**) The position of recently identified residues important for lipid scrambling by nhTMEM16 (red sticks) (Jiang et al., 2017; Lee et al., 2018) are mapped onto the afTMEM16 Ca^2+^-bound (**A**) and Ca^2+^-free (**B**) structures. Critical positions cluster at the rearranging interfaces between helices TM3-6. Of note is V337 (S329 in afTMEM16, shown in yellow) whose mutation to tryptophan increases scrambling activity in the absence of Ca^2+^. In the Ca^2+^-free structure this residue is positioned at the TM4-TM6 contact point, suggesting that this substitution might prevent closure of the pathway.

**Figure 5-Supplement 1.**
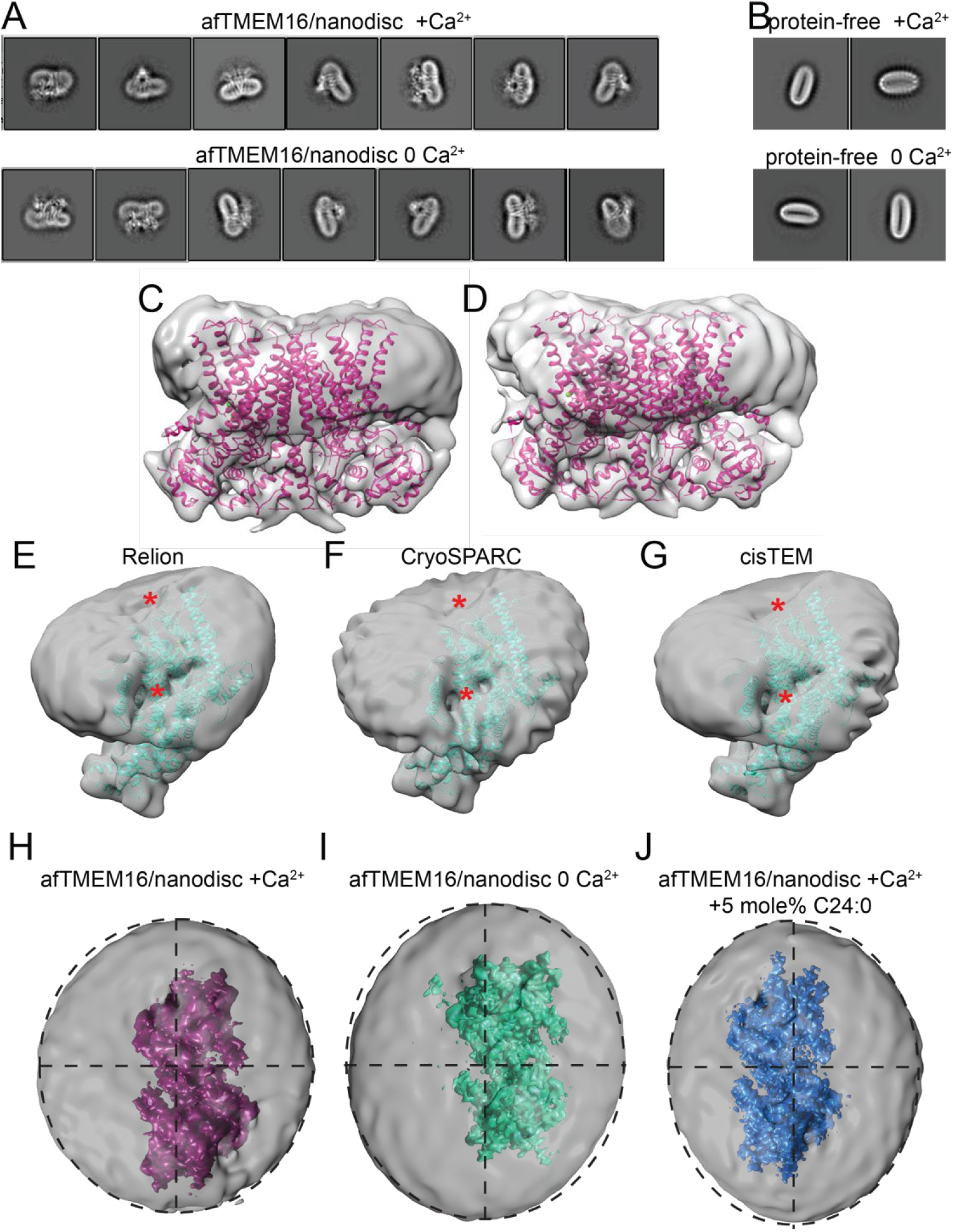
Altered membrane organization induced by activation of afTMEM16. (**A-B**) Close up view of representative 2D classes (box size 275 Å) of Ca^2+^-bound (0.5 mM) (top) and Ca^2+^-free (bottom) nanodisc-reconstituted afTMEM16 (taken from Extended Data Fig. 1 and 3 respectively). Note how the remodeling of the nanodisc induced by afTMEM16 is visible in the 2D classes. (**B**) Close up view of 2D class averages (box size 275 Å) of empty (without protein) nanodiscs from the +Ca2+ (top) and Ca2+-free (bottom) data sets. Note how the empty nanodiscs are flat and not curved. (**C-D**) Unmasked final maps of afTMEM16/nanodisc complex in the presence of 0.5 mM Ca^2+^ low pass filtered to 10 Å from a dataset that resulted in a reconstruction with a final resolution of ~ 8 Å (**C**), or from one of the two data sets used in the final reconstruction that resulted in a protein map with a resolution of 4.5 Å (**D**). Both were processed as described in the methods. (**E-G**) 3D reconstructions from the +Ca^2+^ dataset processed with three different programs for single particle analysis, relion (**E**), cryoSPARC (**F**), and cisTEM (**G**). The membrane bending and reorganization is independent of the program used and is present in all 3D reconstructions. (**H-J**) Top views of the afTMEM16-nanodisc complexes for afTMEM16 +Ca^2+^ (**H,** pink), afTMEM16 Ca^2+^-free (**I**, green), afTMEM16 +Ca^2+^ +5 mole% C24:0 (**J**, blue). The masked final maps are shown inside unmasked maps, which were low pass filtered to 10 Å and are shown at σ=0.25. Dashed circles indicate the oval outline of nanodisc and the dashed lined denote the axis of the oval to highlight the different position of the protein map relative to the nanodisc in the 3 conditions.

**Figure 5-Supplement 2.**
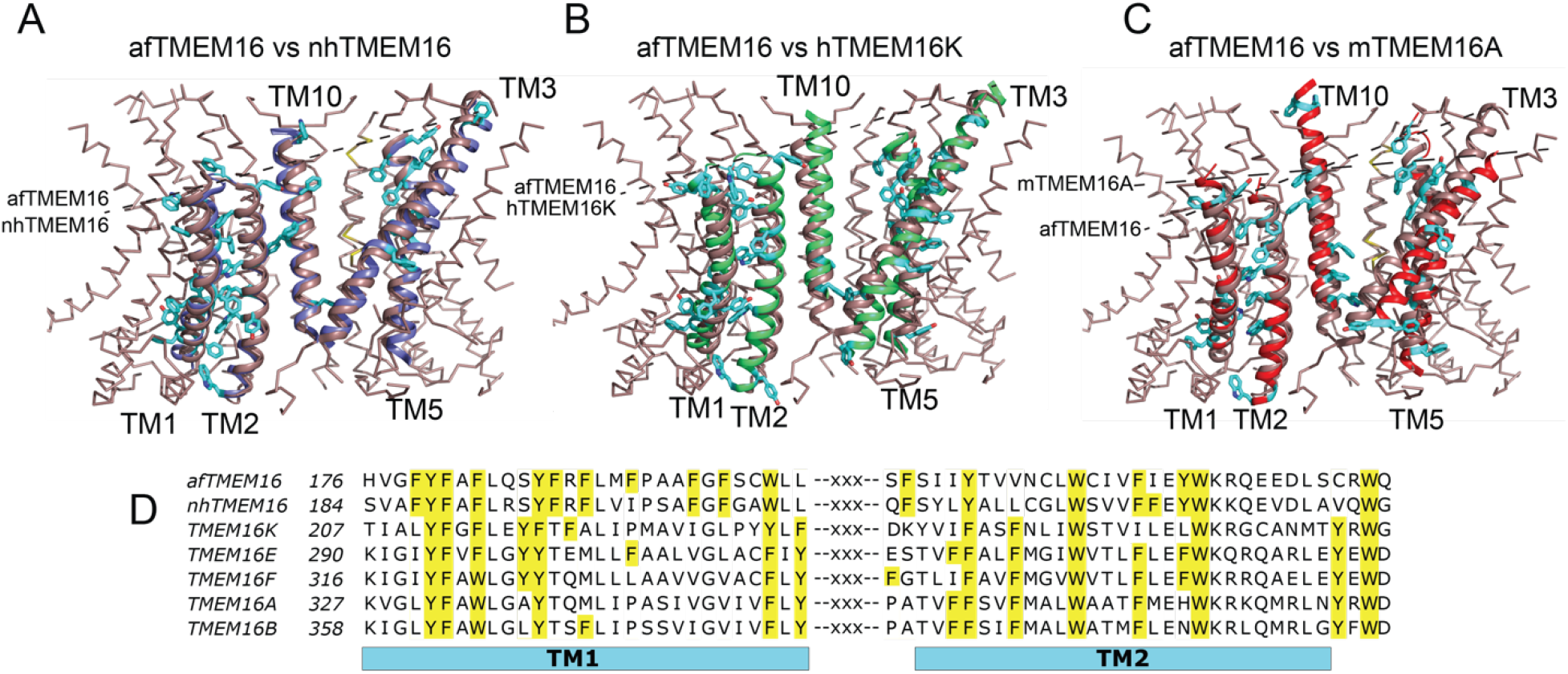
Architecture of the dimer cavity of TMEM16 proteins. (**A-C**) Structural alignment of the dimer cavities of nhTMEM16 (purple) (**A**), hTMEM16K (green) (**B**), or mTMEM16A (red) (**C**) to afTMEM16 (maroon). The transmembrane region of afTMEM16 is shown as ribbon. The five cavity-lining helices (TM1, TM2, and TM10 from one monomer, TM3 and TM5 from the other) from both displayed structures are shown in cartoon representation. Aromatic side chains from nhTMEM16 (**A**), hTMEM16K (**B**), and TMEM16A (**C**) lining the cavity are shown as cyan sticks. The TM1 comprises 29 total residues, of which 11 are aromatic in afTMEM16, 10 in nhTMEM16, 8 in TMEM16K and 7 in TMEM16A. (**D**) Conservation of the aromatic residues in TM1 and TM2 within the TMEM16 family. Sequence alignment was generated using PROMALS3D (Pei and Grishin, 2014). Aromatic residues in TM1 and TM2 are highlighted in yellow.

**Figure 6-Supplement 1.**
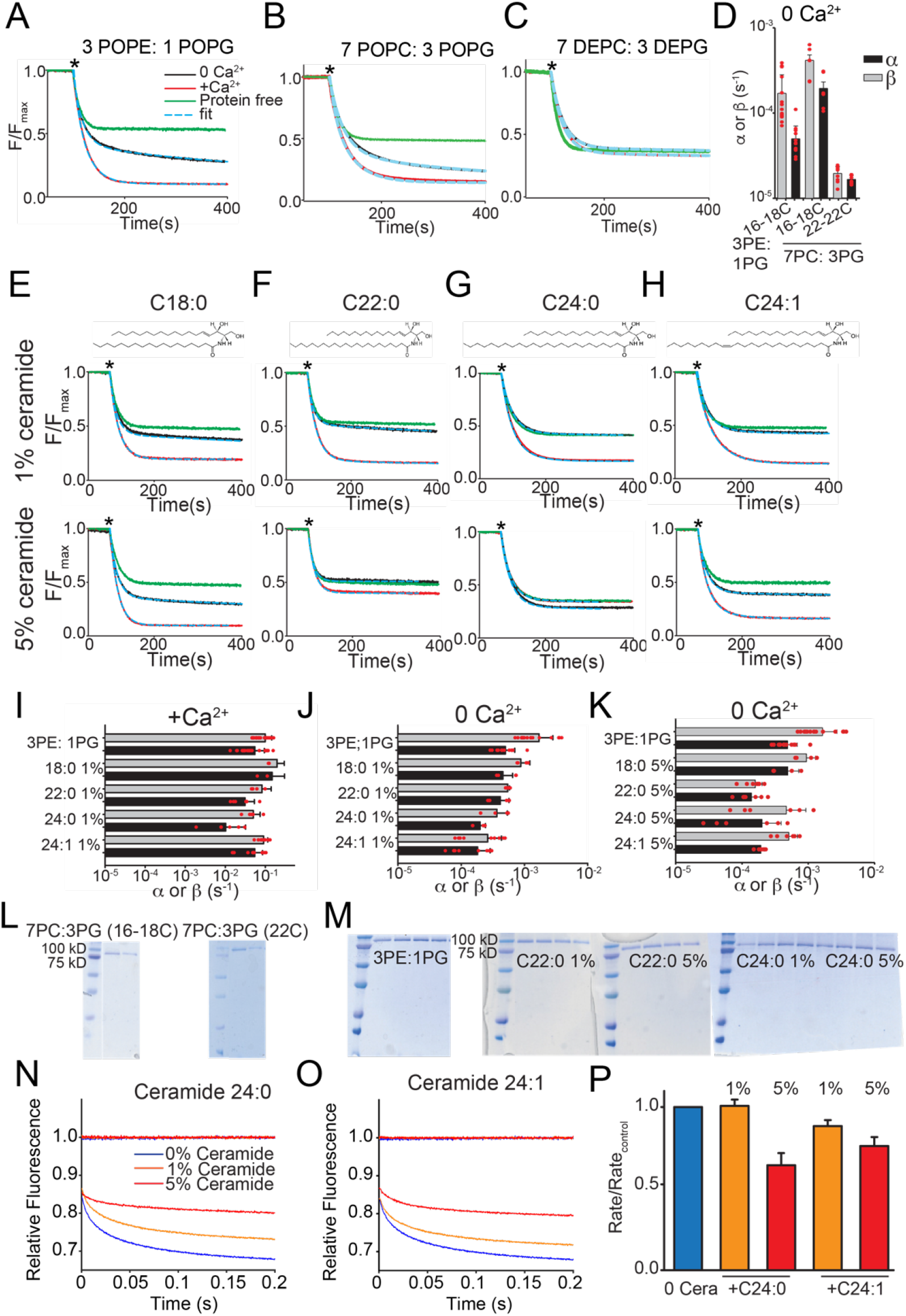
Functional modulation of lipid scrambling by bilayer properties. (**A-C**) Representative time courses of dithionite-induced fluorescence decay in protein-free liposomes (green) or in afTMEM16 proteoliposomes with (red) or without (black) Ca^2+^. Cyan dashed lines represent the fit to Eq. 1 (Malvezzi et al., 2018). * denotes addition of dithionite. Liposomes were formed from 3 POPE: 1 POPG (**A**), 7 POPC: 3 POPG (B) or 7 DEPC: 3 DEPG (**C**) mixtures. (**D**) Forward (α, black) and reverse (β, gray) scrambling rate constants in the absence of Ca^2+^ as a function of lipid chain length. (**E-H**) Representative time courses of dithionite-induced fluorescence decay in control liposomes formed from a 3 POPE: 1 POPG mixtures with Ceramide 18:0 (E), Ceramide 22:0 (**F**), Ceramide 24:0 (**G**) or Ceramide 24:1 (**H**) at 1 mole% (top panels) or at 5 mole% (bottom panels). (**I-K**) Forward (α, black) and reverse (β, grey) scrambling rate constants of afTMEM16 with 1 mole% ceramides with (**I**) or without Ca^2+^ (**J**) or with 5 mole% Ceramide and without Ca^2+^ (**K**). Data in panels (**D)** and (**I-K**) is reported as mean ± St.Dev. Red circles denote individual experiments. Values and exact repeat numbers are reported in Supplementary Tables 2-3. (**L** and **M**) SDS-PAGE gels of liposomes containing afTMEM16 throughout the reconstitution confirming the presence of the protein in the conditions without scrambling activity as compared to the associated control 22C PC-PG (L), C22:0 (M middle), C24:0 (M right). (**L**) Samples were taken from the reconstitution after the first biobead change (left) and after the last (right). (**M**) Samples were taken from the reconstitution after each of the four biobead changes. In all gels the final band represents the protein present in the liposomes when the experiment was completed. (**N-P**) Representative time course of gramicidin A (gA)-mediated Tl^+^-induced quenching of ANTS in 3 POPE:1 POPG large unilamellar vesicles + C24:0 ceramide (L) or C24:1 ceramide (**M**). Blue trace: control–no ceramide; orange trace: 1% ceramide; red trace: 5% ceramide. (**N**) Normalized rate of the Tl^+^-induced fluorescence quenching. The quenching rate was determined by fitting a stretched exponential (Eq. 3) to the fluorescence time course in **L-M**. The quenching rate reflects the gA monomer↔dimer equilibrium that is affected by the chemicophysical properties of the membrane where a reduction in the rates reflects a stiffening and/or reduction in fluidity of the membrane.

**Figure 7-Supplement 1.**
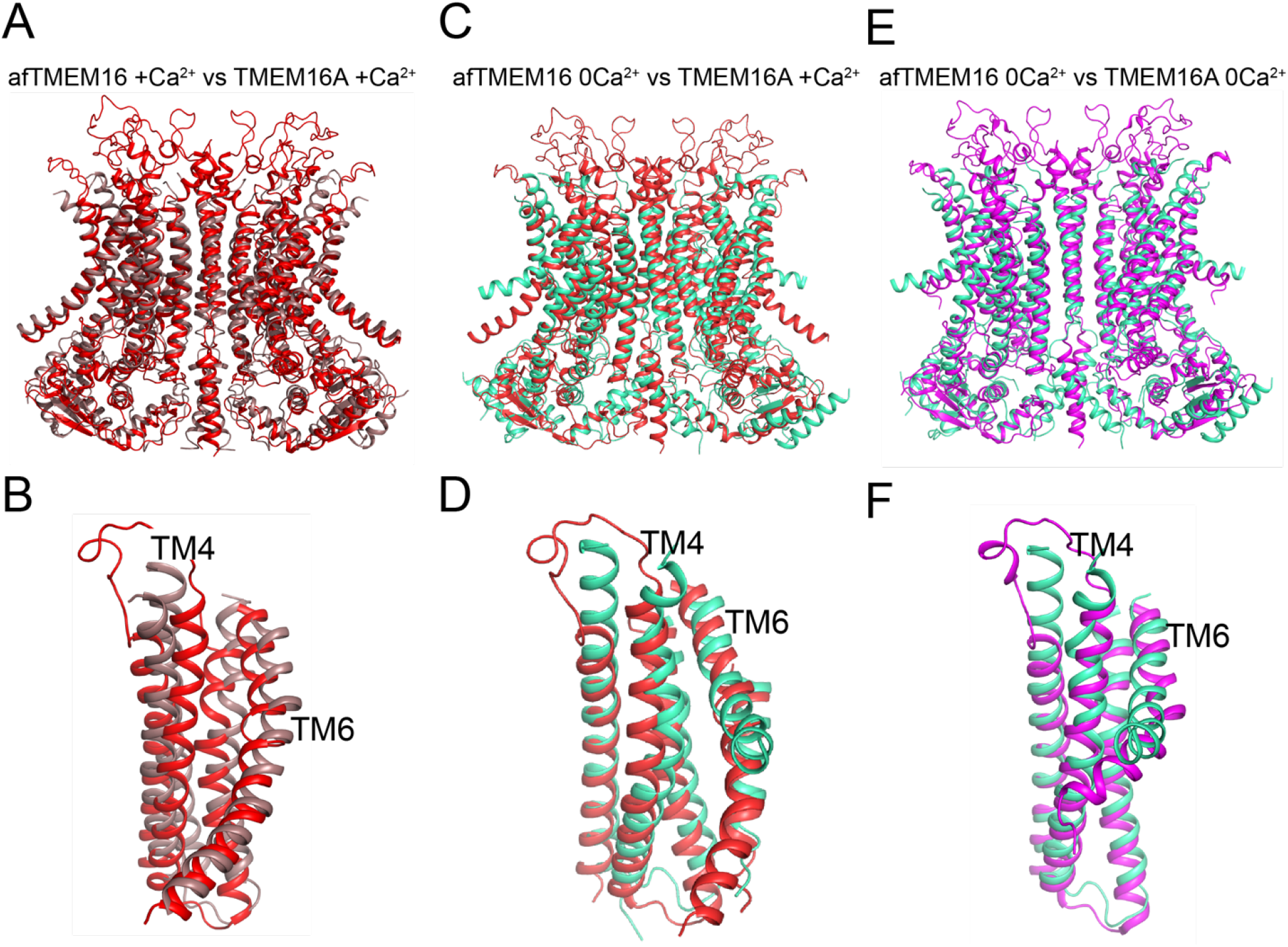
Structural comparison of Ca^2+^-bound and Ca^2+^-free afTMEM16 and TMEM16A. (**A-B**) Alignment of Ca^2+^-bound afTMEM16 (maroon) and Ca^2+^-bound TMEM16A (red, PDBID: 5OYB). A close up view of the pathway is shown in (**B**) to highlight the closure of the permeation pathway in the TMEM16A channel by a rearrangement of TM4. (**C-D**) Alignment of Ca^2+^-free afTMEM16 (cyan) and Ca^2+^-bound TMEM16A (red). A close up view of the pathway is shown in (D) to highlight how the TM4 helices adopt similar conformations. (**E-F**) Alignment of Ca^2+^-free afTMEM16 (cyan) and Ca^2+^-free TMEM16A (magenta, PDBID 5OYG). A close up view of the pathway is shown in (**F**) to highlight how the TM4 helices adopt similar conformations and the TM6 bend at the same points. Note the different orientations of the intracellular portions of the TM6 from the two structures in the absence of Ca^2+^.

